# Single-molecule analysis of PARP1-G-quadruplex interaction

**DOI:** 10.1101/2025.01.06.631587

**Authors:** Paras Gaur, Fletcher E. Bain, Riaz Meah, Maria Spies

## Abstract

The human genome contains numerous repetitive nucleotide sequences that display a propensity to fold into non-canonical DNA structures including G-quadruplexes (G4s). G4s have both positive and negative impacts on various aspects of nucleic acid metabolism including DNA replication, DNA repair and RNA transcription. Poly (ADP-ribose) polymerase (PARP1), an important anticancer drug target, has been recently shown to bind a subset of G4s, and to undergo auto-PARylation. The mechanism of this interaction, however, is poorly understood. Utilizing Mass Photometry (MP) and single-molecule total internal reflection fluorescence microscopy (smTIRFM), we demonstrate that PARP1 dynamically interacts with G4s with a 1:1 stoichiometry. Interaction of a single PARP1 molecule with nicked DNA or DNA containing G4 and a primer-template junction is sufficient to activate robust auto-PARylation resulting in the addition of poly (ADP-ribose) chains with molecular weight of several hundred kDa. Pharmacological PARP inhibitors EB-47, Olaparib and Veliparib differently affect PARP1 retention on G4-containing DNA compared to nicked DNA.

## INTRODUCTION

Poly (ADP-ribose) Polymerase 1 (PARP1) participates in a wide range of cellular processes. It is a key player in genome maintenance and is a universal sensor of DNA damage, which recruits various DNA repair proteins to damaged DNA and catalyzes the addition of poly(ADP-ribose) chains or PARylation (1–8), which is the addition of ADP-ribose (ADPr) units from NAD^+^ to target proteins, forming branched chains of negatively charged poly(ADP-ribose) (PAR) (4,5,9). In its free state PARP1 is composed of several distinct domains arranged like “beads-on-a-string”. Each of these domains has specific functions and interactions. The N-terminal region contains three zinc finger motifs with Zn1 and Zn2 involved in DNA recognition and Zn3 is important for allosteric activation. The BRCT (BRCA1 C-terminal) domain, while dispensable for PARP1 activation, contains auto-modification sites. The WGR (Tryptophan-Glycine-Arginine) domain transfers activation signals from zinc fingers 1 and 2 to the catalytic domain. The HD (helical subdomain) of the catalytic domain serves an auto-inhibitory function. Finally, the ART (ADP-ribosyl transferase) domain contains the enzyme’s active site, and a conserved fold found in all PARP family members (7,10,11). ART domain is composed of a donor (NAD^+^-binding) site that positions the ‘donor’ ADP-ribose for the transferase reaction and an acceptor site that binds either the PARylation target during initiation or the distal ADP-ribose monomer of the growing PAR chain (‘acceptor’) during elongation/branching stages (12). The recognition of exposed bases at the DNA damage site by PARP1 zinc fingers induces PARP1 self-assembly from a “beads-on-a-string” flexible arrangement to a highly organized structure, which requires local unfolding of the HD subdomain (7,8,10,11,13,14). There are several sites of ADP-ribosylation reported in PARP1, specifically D387, E488, and E491 in the BRCT domain and in the flexible linker (15). Additional ADP-ribosylation sites have also been identified in functional domains of PARP1, including all three zinc fingers (16).

Guanine-rich repetitive sequences with the pattern G3+N1-7G3+N1-7G3+N1-7G3+N1-7 in the human genome can fold into non-canonical DNA structures called G-quadruplexes (G4) (17–19). These structures can fold intra and intermolecularly in single-stranded DNA (18,19). Guanines from each G-repeat form a Hoogsteen base pair, creating square coplanar structures called G-quartets which stack on top of each other, stabilized by π-π interactions, with the phosphate backbone forming the corners. The loop sequences between G-repeats extrude outside the structure. G4 stability depends on loop sequence, length, and the number of guanines in the repeats (18–21). The metal cations, particularly K^+^, promote G4 folding and stabilization, while Na^+^ has a moderate and Li^+^ has a minimal effect (22–25). G4s exist in three main topologies: parallel, antiparallel, and hybrid, based on the orientation of the phosphate backbone in the G-repeats (26). G4 structures can stall DNA replication forks (27). During replication, G4 bypass requires rapid recruitment of proteins that can recognize and process these non-canonical DNA structures (28,29).

Recently, several studies have shown that PARP1 interacts with G4 DNA (30,31). G4 binding by PARP1 triggers its catalytic activity and leads to PAR synthesis (21). PARP1 binds with nanomolar affinities to *c-KIT* and *c-MYC* promoter G4 DNA structures but shows little binding to human telomeric G4 DNA (32). In this study we utilized single-molecule techniques to explore PARP1-G4 interaction kinetics, stoichiometry and PARylation.

## MATERIAL AND METHODS

### DNA substrates

DNA oligonucleotides were purchased from Integrated DNA Technologies (Coralville, IA, USA). The sequences of the individual oligonucleotides and additional modifications are listed in **Supplementary Table S1**.

### PARP1 Purification and Fluorescent labeling

Wild type human PARP1 was purified using the protocol published by Pascal lab with minor modifications (33). The concentration of purified PARP1 was calculated by measuring the absorbance at 280 nm, with an extinction of coefficient 120,055 M^-1^cm^-1^.

PARP1 was labeled with Cy3 monofunctional dye (Cytivia, PA23001) or Cy5 monofunctional dye (VWR Catalog# 95017-379). A 16.7x Molar excess of Amersham Cy3 dye or Cy5 dye was added to the PARP1 and incubated for 8 hours at 4°C with rotation. The labeling reaction was stopped with the addition of 50 mM Tris-HCl pH 7.0. The sample was filtered with a 0.22 µM filter and loaded onto a pre-equilibrated Heparin column with a buffer containing 25 mM HEPES-NaOH pH 8.0, 200 mM NaCl, 1 mM EDTA, and 0.1 mM TCEP-HCl. To remove unincorporated dyes, fluorescently labeled PARP1 was eluted from Heparin column with a NaCl gradient ranging from 200 mM to 1M. The PARP1 containing fractions were verified by gel electrophoresis. The labeling efficiency was determined by calculating respective protein and dye concentrations from absorbance at 280 nm (for [PARP1]), 550 nm (for [Cy3]) or 650nm (for [Cy5]) (extinction coefficients: 150,000 M^-1^cm^-1^ for Cy3, 250,000 M^-1^cm^-1^ for Cy5).

### Circular Dichroism

The DNA substrates (see **Supplementary Table 1**) were dissolved in a buffer containing 20 mM Tris, 100 mM KCl, 10 mM MgCl2, and 1 mM EDTA, to achieve a final concentration of 5 μM. The samples were then heated for 5 minutes at 95°C and slowly cooled down to make sure secondary structures were formed. The presence of the DNA secondary structure was probed using JASCO J-810 spectropolarimeter. The measurements were conducted at room temperature, with a scan range of 220 to 320 nm. Each scan consisted of 10 repeats, and the averaged ellipticity values were plotted for each data point.

### Mass Photometry

The Mass Photometry (MP) experiments were conducted on Refeyn Two-MP instrument (Refeyn Ltd., Oxford, UK) on pre-cleaned coverslips (24 mm × 50 mm, Thorlabs Inc., Newton, NJ, USA) with serial washing with deionized water and isopropanol followed by drying. The silicon gaskets (Grace Bio-Labs, Bend, OR, USA) were cleaned in a similar process as coverslips and were placed onto coverslips for the experiments (34). The MP measurements were performed in an MP buffer containing 20 mM Tris pH 7.4, 100 mM KCl, 1 mM EDTA, and 10 mM MgCl2. The calibration was performed using a protein standard mixture: of β-amylase (Sigma-Aldrich, 56, 112, and 224 kDa, St. Louis, MO, USA), and thyroglobulin (Sigma-Aldrich, 670 kDa). Before each experiment, a 15 μL buffer was placed into a chamber formed by coverslip-Gasket and focus was searched and followed by locking it using autofocus function. DNA substrates, PARP1 protein and PARPi [Olaparib, Veliparib and EB-47 (Cat. Nos. 7026, 7579 and 4140, Bio-Techne Tocris)] were added to the chamber and mixed by pipetting. To induce PARylation, 0.2 mM NAD^+^ (#N8535, Sigma-Aldrich) was added. The movies were recorded for 60 s (6000 frames) using AcquireMP (Version 2.3.0; Refeyn Ltd., Oxford, UK). All movies were processed and analyzed using DiscoverMP (Version 2.3.0; Refeyn Ltd., Oxford, UK). Individual molecular weight readings for each experiment were binned into 3 kDa intervals, plotted as histograms, and fitted to multiple Gaussians using GraphPad Prism.

### Single-molecule total internal reflection microscopy

A custom-built prism total internal reflection microscope (TIRFM) was used to perform single-molecule TIRFM experiments (35). The microscope is built on an Olympus IX71 microscope frame and combines 532 nm (Compass 215M-50, Coherent Inc., Santa Clara, CA, USA) and 641 nm (Coherent, Cube 1150205/AD) laser beams using a polarizing beam splitting cube (CVI Melles Griot, PBSH-450-700-050), which are directed at the microscope objective at a 30° angle. TIR is achieved through a UV fused silica pellin–broca prism (325-1206, Eksma Optics, Vilnius, Lithuania) and an uncoated N-BK7 plano–convex lens (LA1213 Thorlabs Inc., Newton, NJ, USA). Photons are collected using a 60X, NA 1.20 water immersion objective (UPLSAPO60XW Olympus Corp., Shinjuku City, Tokyo, Japan), and spurious fluorescent signal is removed using a dual bandpass filter (FF01-577/690-25 Semrock Inc., Rochester, NY, USA). Cy3 and Cy5 emissions are separated using a dual-view housing (DV2 Photometrics, Tucson, AZ, USA) containing a 650 nm longpass filter (T650lpxr Chroma Technology Corp., Bellows Falls, VT, USA), and fluorescent images are captured using an Andor iXon 897 EMCCD (Oxford Instruments, Abingdon, UK).

### Surface tethered DNA single-molecule experiments

Prior to surface tethering, the mixture of biotinylated DNA oligo and indicated G4-forming oligos (see **Supplementary Table 1**) were heated together at 95 °C for 5 min and slowly cooled down to allow for annealing and later diluted to working concentration.

To extend the lifespan of fluorophores in single-molecule experiments, an oxygen scavenging system is necessary to reduce reactive oxygen species (ROS) that cause rapid photobleaching. We utilized 12 mM Trolox (6-hydroxy-2,5,7,8-tetramethylchroman-2-carboxylic acid) and gloxy (catalase and glucose oxidase solution) to reduce ROS effects. Trolox is prepared by adding 60 mg of Trolox powder (238813-5G, Sigma-Aldrich) to 10 mL of water with 60 μL of 2 M NaOH, mixing for 3 days, filtering, and storing at 4 °C. Gloxy is prepared as a mixture of 4 mg/mL catalase (C40-500MG, Sigma-Aldrich) and 100 mg/mL glucose oxidase (G2133-50KU, Sigma-Aldrich) in wash buffer (25 mM Tris-HCl pH 7.0, 140 mM KCl); special KCl of Spectroscopy grade (#39795, Alfa Aesar, Haverhill, MA, USA) was used for all single molecule experiments.

The method for conducting single-molecule TIRFM PARP1-DNA binding experiments is as follows and includes the DNA substrates listed (**Supplementary Table 1**). Prior to the experiment, quartz slides (25 mm × 75 mm × 1 mm #1x3x1MM, G. Frinkenbeiner, Inc., Waltham, MA, USA) and cover glass (24 mm × 60 mm-1.5, Fisherbrand, Fisher Scientific, Hampton, NH, USA) are treated with passivation and PEGylation (36). The flow cell is treated with 0.2 mg/mL NeutrAvidin (#3100, Thermo Fisher Scientific, Waltham, MA, USA) to tether biotinylated molecules to the flow cell surface. Excess neutravidin is removed with flow of 1 mL wash buffer. To prepare the flow cell for imaging, 100 pM of biotinylated-DNA substrates are added and incubated for 3 min. Excess DNA is then removed using a wash buffer. For image collection, imaging buffer (containing 25 mM Tris-HCl pH 7.0, 140 mM KCl, 10 mM MgCl2, 1 mg/mL BSA, 1 mM DTT, 0.8% *w/v* D-glucose, 12 µM glucose oxidase, 0.04 mg/mL catalase and Trolox) is added to the flow cell. The images are then collected using custom software, single.exe (generously provided by the Taekjip Ha Lab, Harvard Medical School), with 532 nm laser power (for Cy3) and 640 nm laser power (for Cy5) were set to 45 mW. Image collection begins using 100 ms time resolution, gain of 290, background set to 400 and correction set to 1200. The range of concentration in pM of Cy3-PARP1 (Cy5-PARP1) is added to the flow cell with or without Olaparib, Veliparib and EB-47 (Cat. Nos. 7026, 7579 and 4140, Bio-Techne Tocris). Images are collected for a total of 4000 or 6000 frames (400 or 600 s).

### Single-molecule data analysis

Fluorescent spot finding and trajectory extraction are done using an IDL script (generously provided by the Taekjip Ha Lab, Harvard Medical School). Individual trajectories are then chosen for analysis using in-house MATLAB scripts. Trajectories were selected based upon the following criteria: no fluorescent intensity changes are present prior to frame 300 (in case of NAD^+^ experiments), baseline must be consistent throughout the trajectory and 2 fluorescent events persisting above the baseline for 3 frames must be present. The selected trajectories are then imported into hFRET (37) and fit to different states of fluorescent intensity. The best fit was determined from the largest log evidence, a lower bound comparison of the three tested models. The dwell times for the events in each state are then extracted using KERA MATLAB software (38), binned, and plotted as a histogram. The histograms were then fit to a single or double exponential decay using GraphPad Prism. The best fit was determined from an F-test comparing the two exponential fits. For all cases where a single exponential decay was the best fit, we obtained a single dissociation rate constant koff, which was independent of the protein concentration. The dwell-time distribution constructed from all unbound data and fitted with a single-exponential function yielded the association rate von, which increases with increasing protein concentration. The association rate constants kon were calculated from respective von values and protein concentrations adjusted by labeling efficiency (85% and 100% labeling efficiency for Cy3-PARP1 and Cy5-PARP1). The equilibrium dissociation constant was calculated from the rate constants as Kd = koff/kon (Table 1). In one case (nicked DNA substrate with EB-47) the “on” dwell time distribution was best fit to a double exponential function yielding koff (Fast) and koff (Slow). To calculate the respective Kd, the protein concentration was adjusted based on Fast and Slow contributions to the double exponential function.

**Table 1.**

| **Protein-DNA** | **Inhibitor** | **Protein** | **State 1 ($k_{on}$) \***<br>**$s^{-1} M^{-1}$** | **State 2 ($k_{off}$)**<br>**$s^{-1}$** | **State 2 ($\tau$)**<br>****s**** | **Kd = $k_{off}/k_{on}$**<br>**(nM)** |
| --- | --- | --- | --- | --- | --- | --- |
| PARP1-nicked | N/A | 100 pM | $(1.41 \pm 0.24) \times 10^8$ | $0.56 \pm 0.03$ | $1.79 \pm 0.09$ | $3.96 \pm 0.2$ |
| PARP1-nicked | Veliparib (75 nM) | 100 pM | $(2.82 \pm 0.24) \times 10^8$ | $0.75 \pm 0.05$ | $1.33 \pm 0.08$ | $2.67 \pm 0.16$ |
| PARP1-nicked | Olaparib (30 nM) | 100 pM | $(4.35 \pm 0.35) \times 10^8$ | $0.36 \pm 0.02$ | $2.78 \pm 0.12$ | $0.82 \pm 0.04$ |
| PARP1-nicked | EB-47 (75 nM) | 100 pM | **Fast**<br>$(2.02 \pm 0.02) \times 10^9$<br>**Slow**<br>$(9.99 \pm 0.98) \times 10^9$ | **Fast**<br>$0.85 \pm 0.07$<br>**Slow**<br>$0.13 \pm 0.01$<br>**Percent Fast**<br>83.15<br>**Percent Slow**<br>16.85 | **Fast**<br>$1.18 \pm 0.1$<br>**Slow**<br>$7.72 \pm 0.53$ | **Fast**<br>$0.42 \pm 0.04$<br>**Slow**<br>$0.01 \pm 0$ |
| PARP1-cKITG4 | N/A | 200 pM | $(5.735 \pm 0.141) \times 10^9$ | $0.25 \pm 0.01$ | $3.95 \pm 0.22$ | $0.04 \pm 0.002$ |
| PARP1-cKITG4 | Veliparib (75 nM) | 200 pM | $(2.265 \pm 0.112) \times 10^9$ | $0.35 \pm 0.02$ | $2.87 \pm 0.16$ | $0.15 \pm 0.01$ |
| PARP1-cKITG4 | Olaparib (30 nM) | 200 pM | $(1.147 \pm 0.088) \times 10^9$ | $0.31 \pm 0.01$ | $3.24 \pm 0.07$ | $0.27 \pm 0.01$ |
| PARP1-cKITG4 | EB-47 (75 nM) | 200 pM | $(2.824 \pm 0.294) \times 10^8$ | $0.34 \pm 0.01$ | $2.95 \pm 0.09$ | $1.2 \pm 0.04$ |
| PARP1-cKITG4-PT | N/A | 100 pM | $(2.35 \pm 0.12) \times 10^8$ | $0.76 \pm 0.02$ | $1.31 \pm 0.04$ | $3.24 \pm 0.09$ |
| PARP1-cKITG4-PT | Veliparib (75 nM) | 100 pM | $(2.59 \pm 0.353) \times 10^8$ | $0.99 \pm 0.04$ | $1.01 \pm 0.04$ | $3.82 \pm 0.14$ |
| PARP1-cKITG4-PT | Olaparib (30 nM) | 100 pM | $(2.12 \pm 0.12) \times 10^8$ | $0.84 \pm 0.02$ | $1.19 \pm 0.03$ | $3.96 \pm 0.11$ |
| PARP1-cKITG4-PT | EB-47 (75 nM) | 100 pM | $(2.12 \pm 0.47) \times 10^8$ | $0.61 \pm 0.02$ | $1.65 \pm 0.06$ | $2.87 \pm 0.1$ |

## RESULTS

### Design of the DNA substrates for Single-molecule total internal reflection microscopy and mass photometry studies

In this study we utilized three main DNA substrates designated as cKITG4 (G-quadruplex forming sequence that is present in promoter region of the c-KIT proto-oncogene (39)), nicked (single strand break) and cKIT G4 with a primer-template junction (cKITG4-PT). The cKITG4-PT consists of cKITG4 flacked by either two hairpins or a hairpin and a biotinylated duplex and is designed to resemble a DNA structure appearing in cells when the DNA replication is stalled by the G4. It has elements of both dsDNA and G4. All substrates used in the mass photometry (MP) studies comprised single oligonucleotides to prevent the appearance of open dsDNA ends. The nicked and cKITG4-PT substrates used in the single-molecule total internal reflection fluorescence microscopy (smTIRFM) studies were assembled using two oligonucleotides with the free end being protected by the hairpin, while the biotin-containing end used for surface-tethering being protected by the surface. The simple cKITG4 substrate in smTIRFM analyses was biotinylated at the 3’ end (**Supplementary Table 1**).

Before starting single molecule experiments, we verified folding of the DNA substrates using CD spectroscopy (**Supplementary Figure 1**). The simple cKITG4 yielded an expected spectrum with a negative peak around 242 nm and a positive peak at 262 nm (40). The nicked substrate produced a spectrum characteristic of dsDNA (41), and cKITG4-PT spectrum was the sum of that for cKITG4 and nicked substrates (**Supplementary Figure 1A and B**).

### Dynamic interaction of PARP1 with surface tethered DNA

We first used smTIRFM to study the dynamic interactions of PARP1 protein and surface-tethered DNA, and to determine binding stoichiometry.

The biotinylated DNA substrates (**Supplementary Table 1**) were folded and annealed in KCl. The folded DNA substrates (100 pM) were immobilized on the surface of the TIRFM reaction chamber, and Cy3-labeled PARP1 (85% labeling efficiency) was flowed into the chamber in the presence of K^+^ buffer (**Table 1** lists [Cy3-PARP1] in each experiment). The movies were recorded for 6000s yielding fluorescence trajectories that show the time-based changes in Cy3 fluorescence in a specific location in the TIRFM reaction chamber and represent Cy3-PARP1 binding to and dissociating from individual surface-tethered DNA molecules (**Figure 1 and Supplementary Figure 2-3**). We observed fluctuations in the magnitude of Cy3-labeled PARP1 fluorescence when bound to DNA, which could be interpreted as either the presence of multiple PARP1 molecules binding or as photophysical effects. Notably, N-terminal Cy3 labeling of PARP1 can position the Cy3 fluorophore in such a way that PARP1 binding to DNA affects the Cy3 environment resulting in protein enhanced fluorescence (PIFE). This contrasts with a study where a Halo tag was placed on the N-terminus for Cy3 labeling, resulting in a less noisy signal (42). To resolve whether multiple PARP1 molecules bind simultaneously to the surface-tethered DNA substrate, a 1:1 mixture of PARP1 proteins labeled with either Cy3 or Cy5 (100 % labeling efficiency) fluorescence dye was utilized for two-color smTIRFM experiments (**Figure 1 A and B**). Prior to smTIRFM analysis, the DNA binding capacity and enzymatic activity of Cy3 and Cy5 PARP1 for PARylation were verified in MP experiments and determined to be identical to that of the unlabeled protein (**Supplementary Figure 5**). Within the smTIRFM reaction chamber, 100 pM of cKITG4 DNA was immobilized, followed by the infusion of equimolar concentrations of Cy3- and Cy5-labeled PARP1 (100 pM). Our observations of individual molecule trajectories revealed an absence of simultaneous two-color fluorescence events. We consistently observed either red or green signals intermittently appearing in the same trajectory, indicating that only one PARP1 molecule occupied the cKITG4 at a given time (**Figure 1A**). This observation confirms that only a single PARP1 molecule can simultaneously bind to the G4.

**Figure 1.**
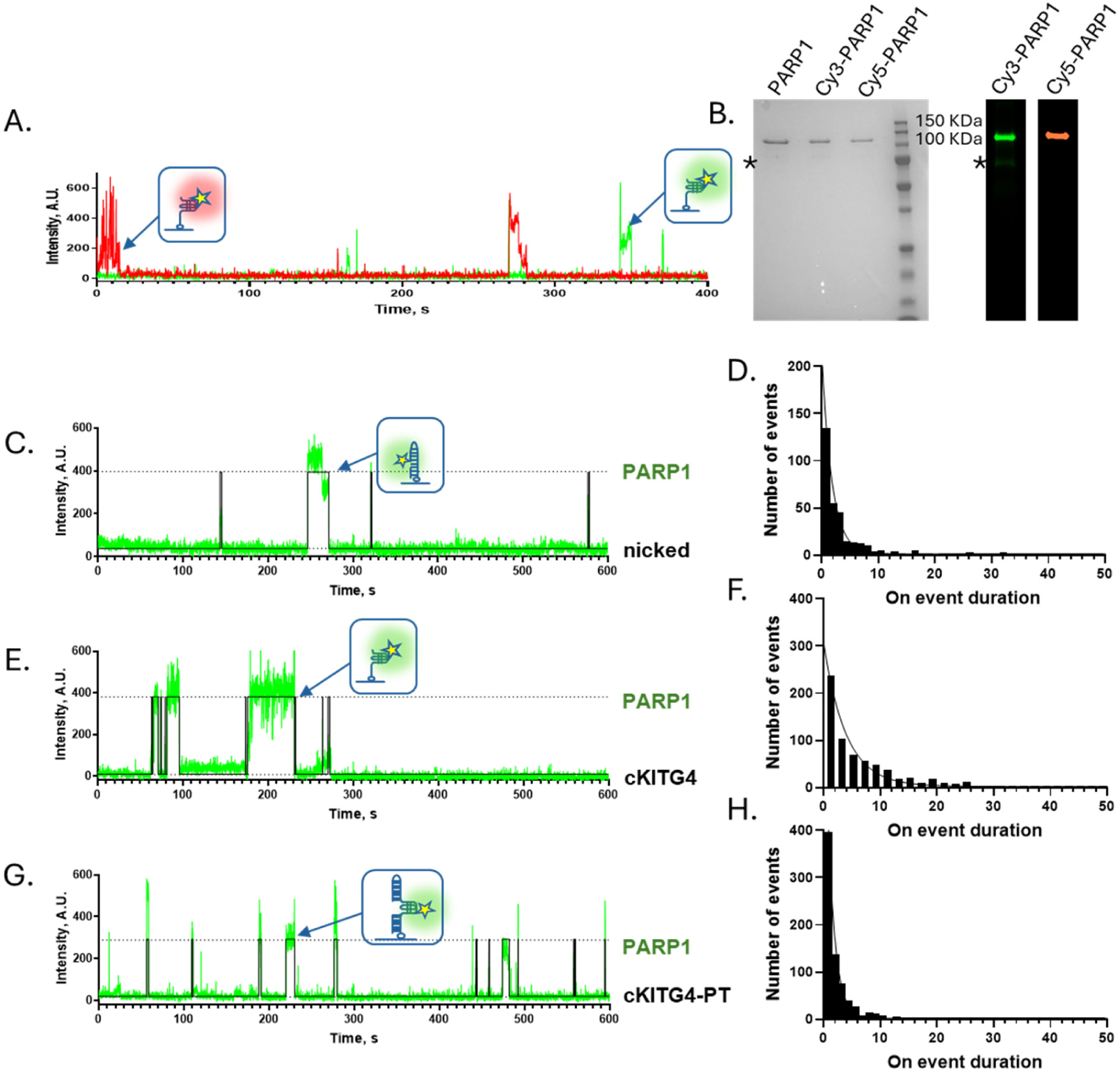
Dynamic binding of PARP1 monomers to the surface-tethered DNA substrates. **A.** Raw smTIRFM fluorescence trajectory showing Cy3 and Cy5-PARP1 infused in single molecule slide chamber and red and green fluorescence signal suggest Cy5-PARP1 and Cy3-PARP1 binding on cKIT DNA**. B.** SDS-PAGE gel of purified PARP1 stained with Coomassie Brilliant Blue and the same gel with fluorescence scan showing Cy3 and Cy5 labeled PARP1. **C, E, and G.** Representative trajectories with nicked, cKIT and cKIT-PT DNA substrate tethered to the surface and Cy3-PARP1 being infused in the chamber. The raw fluorescence data are shown in green, overlaid with an idealized trajectory represented by a black line. **D, F, and H.** Dwell-time distributions for nicked, cKIT and cKIT-PT DNA substrates which were constructed from the “on” dwell times and fitted using a single-exponential function. Note: Asterisk (*) indicates a consistent PARP1 contaminant.

After confirming the binding stoichiometry, the data with cKITG4, nicked and cKITG4-PT were fitted with 2 state model using hFRET (37). Note that the two-state model was also a preferred model selected by hFRET for data collected for all substrates. In the representative trajectories, raw fluorescence data shown in green are overlaid with an idealized trajectory represented by a black line. We observed dynamic binding and dissociation of PARP1 protein to/from each DNA substrate with dwell times lasting a few seconds (**Figure 1 and Supplementary Figures 3-4**).

No binding was observed in control experiments that had no surface-tethered DNA. All trajectories in each experiment were collectively analyzed using hFRET (37) and KERA (38) as discussed in the Materials and Methods section. The 2 states are labeled as “off” (unbound/free DNA) and “on” (bound). The dwell time is represented by Tau (τ), and we obtained values for both the “on” dwell time (τon) and “off” dwell time (τoff) for each data set.

We observed a dwell time of 1.79 ± 0.09 s on nicked DNA, a canonical PARP1 substrate. In contrast, cKITG4 DNA substrates exhibited a longer dwell time of 3.95 ± 0.22 s, indicating a 2.2-fold increase in residence time when bound to G-quadruplexes. Furthermore, our analysis of cKITG4-PT substrate revealed a dwell time of 1.31 ± 0.04 s, suggesting distinct behavior compared to both nicked and cKITG4 substrates. Equilibrium dissociation constants calculated for nicked and cKITG4 substrates were consistent with a trend observed for “on” dwell times, with nicked-PARP1 exhibiting a Kd of 3.96 ± 0.20 nM and cKITG4-PARP1 displaying a Kd of 0.04 ± 0.002 nM, a 99-fold difference suggesting higher affinity for G4 DNA. In contrast, the Kd calculated for cKITG4-PT substrate 3.24 ± 0.09 nM, similar to the nicked DNA.

To further explore PARP1 interactions with G4 substrates, we investigated their binding to other G4-forming DNA substrates such as cMYCG4 (G4 forming sequence located in the promoter region of cMYC) and hTELG4 (G4 forming sequence present in the ends of human telomeres). While cMYCG4 exhibited binding behavior like cKITG4, interactions with hTELG4 were considerably weaker, confirming a potential role of G4 topology in these interactions (**Supplementary Figure 6**). Additionally, we examined PARP1 binding to non-G4 structures, including double-stranded DNA (dsDNA) and single-stranded DNA (ssDNA) (**Supplementary Figure 7**). Given PARP1’s role as a DNA damage sensor, it was not surprising to observe robust interaction with dsDNA and weaker binding to ssDNA in the smTIRFM experiments.

### PARP1 interacts with G4-containing DNA with a 1:1 stoichiometry

A DNA substrate that PARP1 would encounter in the cell at the sites of stalled DNA replication contains a folded G4 and a primer-template junction, which is also one element that can be found at PARP1’s canonical nicked DNA substrates. To investigate how many PARP1 molecules can simultaneously bind G4 DNA, we utilized MP, a single-molecule technique that uses interferometric light scattering to accurately measure masses of macromolecules and macromolecular complexes as they transition from the solution to the glass surface of the MP slide (43,44).

Full-length human PARP1 protein was expressed and purified using a bacterial expression system (**Figure 1B**) and we noticed consistent impurity at around ∼70 kDa size which we have marked with an asterisk (**Figure 2**), and used as an internal control for MP experiments.

**Figure 2.**
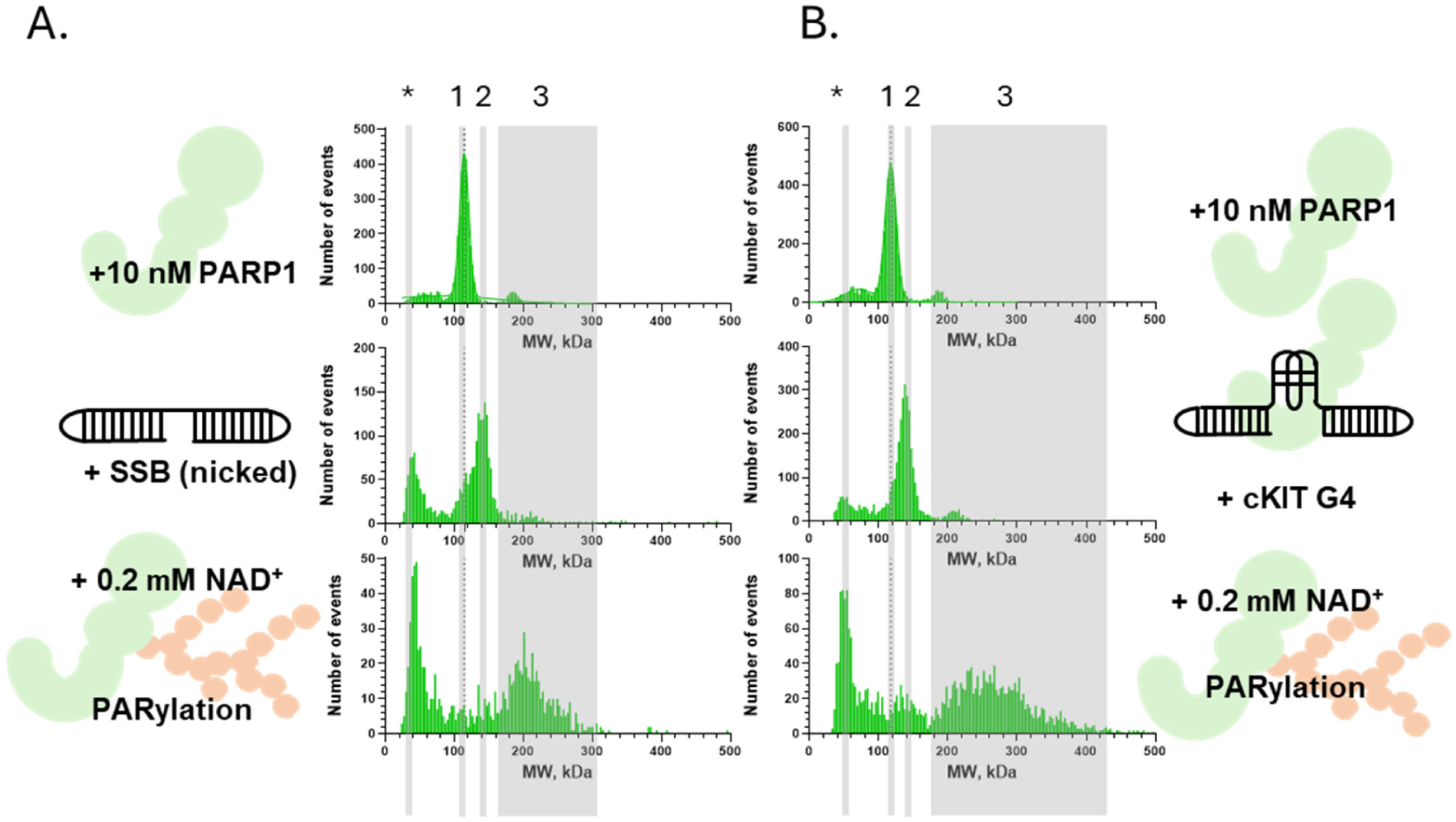
Mass Photometry Analysis of the PARP1 Enzymatic Activity. **A.** PARP1 binding to and activity on nicked DNA substrate: Peak 1 represents PARP1 alone. Following the addition of nicked DNA, Peak 2 emerges showing the PARP1-nicked complex, and after the addition of NAD⁺, Peak 3 forms corresponding to PARylated PARP1, indicating successful PARylation. **B.** PARP1 activity with cKITG4 substrate: Peak 1 represents PARP1 alone, Peak 2 corresponds to PARP1 in complex with cKITG4, and Peak 3 shows robust PARylation. Note: Asterisk (*) indicates a consistent PARP1 contaminant that serves as internal control.

We used nicked DNA (∼22 kDa) and cKITG4-PT (∼39 kDa) substrate to study their complexation with PARP1 protein (113 kDa). Note that because PARP1 can efficiently bind dsDNA ends, all our substrates were annealed from single oligonucleotides (**See Supplementary Table 1**). Initially, 10 nM of PARP1 protein was introduced to MP, showing the expected peak at ∼113 kDa (along with the ∼70 kDa impurity). Subsequently, 10 nM nicked DNA was added, and we observed the expected size shift of 22 kDa, with a peak at ∼135 kDa. Similarly, when cKITG4-PT-nicked substrate was introduced, we observed a size shift from 113 kDa to ∼152 kDa with the 70 kDa impurity remaining in the same spot, indicating that the impurity is not involved in DNA binding. With both DNA substrates, we were able to verify the interaction with PARP1 protein. Different concentrations of DNA and/or proteins were introduced on the MP glass slide in combination, and we observed shifts of 1 molecule of PARP1 and 1 molecule DNA in each case, suggesting 1:1 complex (**Figure 2 A and B**). In the case of cKITG4 DNA substrate without dsDNA feature, we did not observe visible shift in molecular weight likely due to the small size of the DNA substrate (**Supplementary Figure 8**), as we readily observe binding in the smTIRFM experiments. Additionally, as a control, we tested PARP1 complexation with ssDNA, and didn’t observe any binding and subsequent PARylation upon addition of NAD^+^ (**Supplementary Figure 9**).

### A monomer of PARP1 bound to nicked or G-quadruplex containing DNA is sufficient to activate a robust self-PARylation

To induce PARylation in the MP experiments, we introduced 200 μM NAD^+^ to equimolar concentrations (10 nM each) of PARP1 and either nicked DNA or cKITG4-PT DNA. This resulted in a robust PARylation, which manifests as a new higher molecular weight peak (**Figure 2A** and **B**). Time-based measurements demonstrated that PARP1-mediated PARylation saturated within approximately 4 minutes (**Supplementary Figure 10**), reaching a maximum size of ∼350 kDa and ∼500 kDa for nicked and cKITG4-PT DNA, respectively. Notably, PARylation was significantly more pronounced with cKITG4-PT DNA compared to nicked DNA. Based on the observed size shifts of the complex, it is estimated that approximately 503 and 815 ADP-ribose units were attached to PARP1 in the case of nicked and cKITG4-PT DNA, respectively.

### PARP1 inhibitors do not prevent PARP1-G4 DNA interaction

PARP inhibitors (PARPi) exhibit a spectrum of PARP1 trapping activity, by either increasing or decreasing the PARP1 residence time on nicked DNA (13,42). It is unknown whether the same effect can be observed on the G4-containing DNA substrates. We studied three PARPi, which have been previously characterized for their proretention/prorelease activities on nicked DNA; Olaparib, an FDA-approved drug for treatment of recombination deficient cancers (45,46), veliparib (47), another cancer drug which is under clinical trials, and EB-47 a non-clinical PARP inhibitor (48). It was reported that Olaparib moderately increases PARP1 residence on nicked DNA, while EB-47 significantly increases, and Veliparib promotes dissociation/release of the PARP1-nicked DNA complex (13,42). In the MP experiments, 10 nM PARP1 was incubated with Olaparib, Veliparib, or EB-47 at indicated concentrations and then combined with 10 nM of nicked or cKITG4-PT DNA. While none of the three inhibitors interfered with PARP1 binding to either DNA substrate, they all inhibited the PARylation (**Figure 3** and **Supplementary Figure11**).

**Figure 3.**
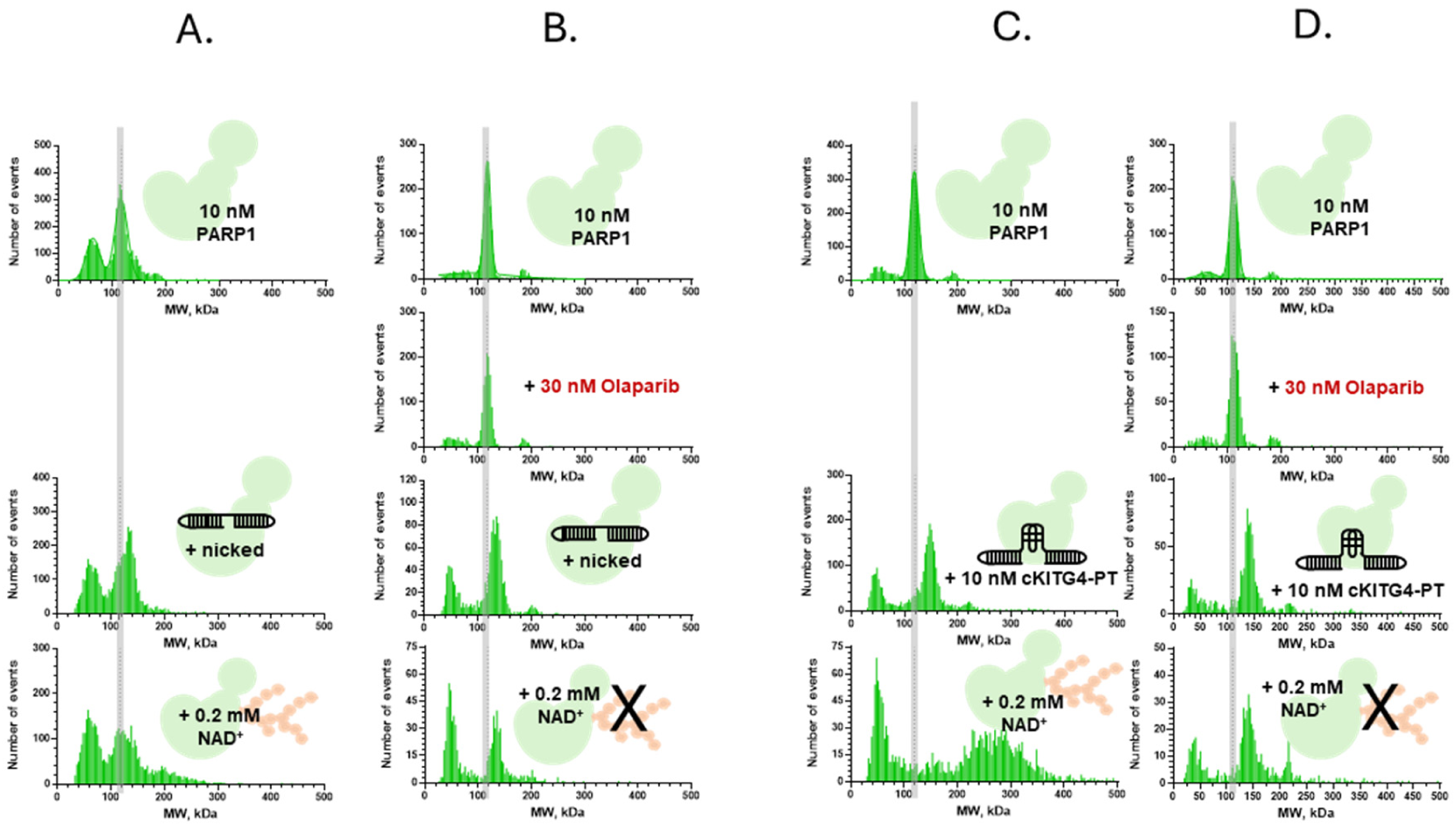
Mass photometry analysis of DNA substrate interactions with PARP1 in presence of PARPi. Panels A and C depict nicked and cKIT-PT DNA substrates, revealing a molecular weight shift (indicated by the transition from a gray line) upon PARP1 addition. This shift suggests the formation of PARP1-nicked and PARP1-cKIT-PT complexes. Following the introduction of 0.2 mM NAD^+^, there is a clear formation of high molecular weight PAR chains, demonstrating the PARylation process. In contrast, panels B and D illustrate the same experimental setup but with the addition of PARPi (Olaparib), a PARP inhibitor. In this scenario, upon NAD+ addition, no PAR chain formation is observed, highlighting the inhibitory effect of Olaparib on PARP1’s enzymatic activity.

In smTIRFM experiments that followed PARP1 binding to nicked DNA we observed a similar trend as previously reported, but different effect of the inhibitors on PARP1 interaction with G4-containing substrates. In these experiments, the biotinylated nicked, cKITG4 and cKITG4-PT DNA substrates were immobilized on a reaction chamber surface, and Cy3-labeled PARP1 molecules were infused in the presence of inhibitors (75 nM Veliparib, 30 nM Olaparib and 75 nM EB-47). **Figure 4** and **Supplementary Figures 12-13** show respective dwell time distributions, **Table 1** lists calculated rate and equilibrium constants, while representative trajectories are shown in **Supplementary Figures 14-22**.

**Figure 4.**
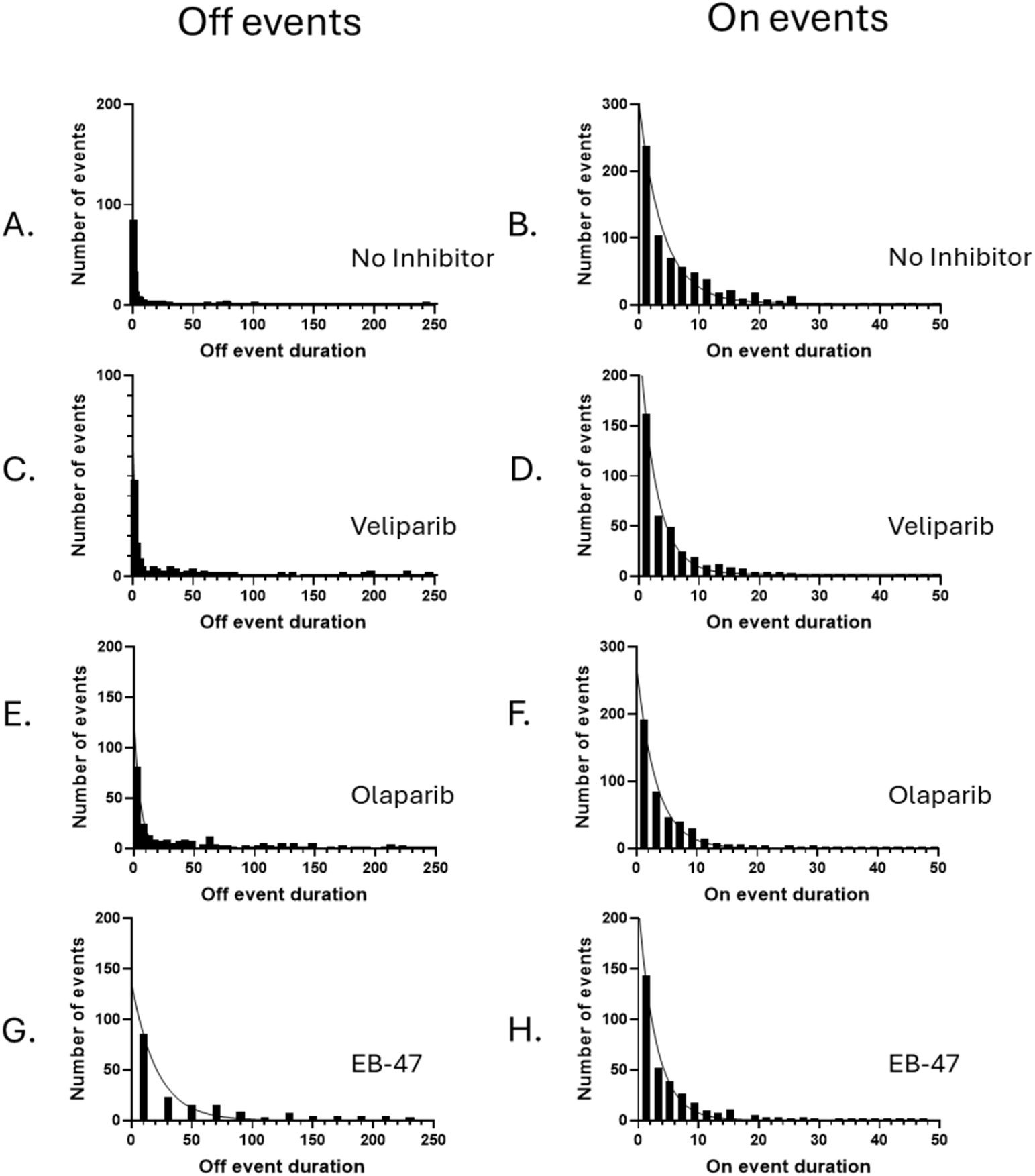
Dwell time distribution of PARP1 on cKITG4 DNA in the presence of PARPi. Biotinylated partial duplex DNA containing the cKIT sequence was immobilized on a surface, while Cy3-labeled PARP1 (Cy3-PARP1) was infused into the reaction chamber with or without PARPi. Representative fluorescence trajectories (green) overlaid with idealized fits (black) are shown in Supplementary Figure 14-22. Dwell-time distributions, constructed from all “OFF” and “ON” states, were analyzed and fitted using a single-exponential function. **(A-B)** Cy3-PARP1 in the absence of PARPi. **(C-D)** Cy3-PARP1 in the presence of Veliparib. **(E-F)** Cy3-PARP1 in the presence of Olaparib. **(G-H)** Cy3-PARP1 in the presence of EB-47.

For nicked DNA, dwell time constants (τ) for PARP1 with Veliparib and Olaparib were 1.33 ± 0.08 s and 2.78 ± 0.12 s, respectively. Addition of EB-47 changed the “on” dwell time distribution of PARP1/cKITG4-PT to a double exponential decay with τfast of 1.18 ± 0.1 s and τslow of 7.72 ± 0.5 s. All other “on” and “off” dwell time distributions were best fit with respective single exponential functions. The preference of a double exponential fit in the case of cKITG4-PT suggests two distinct complexes with different stabilities. Corresponding equilibrium dissociation constants were 2.67 ± 0.20 nM for PARP1 with Veliparib, and 0.82 ± 0.04 nM for Olaparib and for EB-47 as Kdfast 0.42 ± 0.04 and Kdslow 0.01 ± <0.01.

It is important to note that while PARPi modulated PARP1 Kds for the respective DNA substrates, all these affinities were in a low nM range and that the MP experiments, therefore, were conducted under the stoichiometric binding conditions.

PARPi increased dwell times for nicked DNA compared to PARP1 alone (1.79 ± 0.09 s), in particular Olaparib and EB-47 increased dwell times by ∼1.55x and ∼4.3x and veliparib decreased the dwell time ∼0.74x respectively. The trend we observed is similar to a study which reported change in relative retention efficiency in single-molecule colocalization assay as ∼-8% change in PARP1-nicked DNA retention time with Veliparib, ∼7% with Olaparib and ∼15% with EB-47. Increase in PARP1 retention for EB-47 (∼15%) and a modest increase for Olaparib, whereas Veliparib induced a decrease in retention follows a similar pattern to our smTIRFM studies (see Table 1) (42). These results demonstrated that EB-47 and Olaparib modify the PARP1-DNA complex increasing the retention, whereas veliparib facilitated the release.

However, with cKITG4 DNA, dwell times did not exhibit a clear trend with PARPi but were somewhat reduced compared to uninhibited PARP1 (see **Table 1**). The cKITG4-PT substrate showed no significant changes in dwell times or dissociation constants with or without PARPis in smTIRFM.

### NAD^+^ Reduces PARP1 binding to DNA

Our MP experiments revealed a 1:1 interaction between PARP1 molecules and cKITG4-PT DNA, accompanied by robust PARylation (**Figure 2B**). To investigate whether PARylated PARP1 retains the ability to interact with DNA substrates, we carried out smTIRFM experiments in the presence of NAD^+^, a substrate for PARP1 enzymatic activity (49–53). The presence of NAD^+^ in the DNA binding experiments creates an environment that promotes auto-PARylation of PARP1. The addition of 5 mM NAD^+^ with PARP1 to smTIRFM chamber resulted in two distinct changes in PARP1-DNA complexes. First, the PARP1 association to DNA was dramatically reduced. Single-molecule trajectories from these experiments were largely devoid of more than one binding event (**Figure 5**), further demonstrating the loss of DNA binding. Notably, binding events are more frequently observed close to the beginning of the trajectories, with over two-fold higher frequency of binding in the first 300 seconds of the experiments compared to the last 300 seconds. This suggests that the initial PARP1 binding triggers autoPARylation, which in turn precludes subsequent binding.

**Figure 5.**
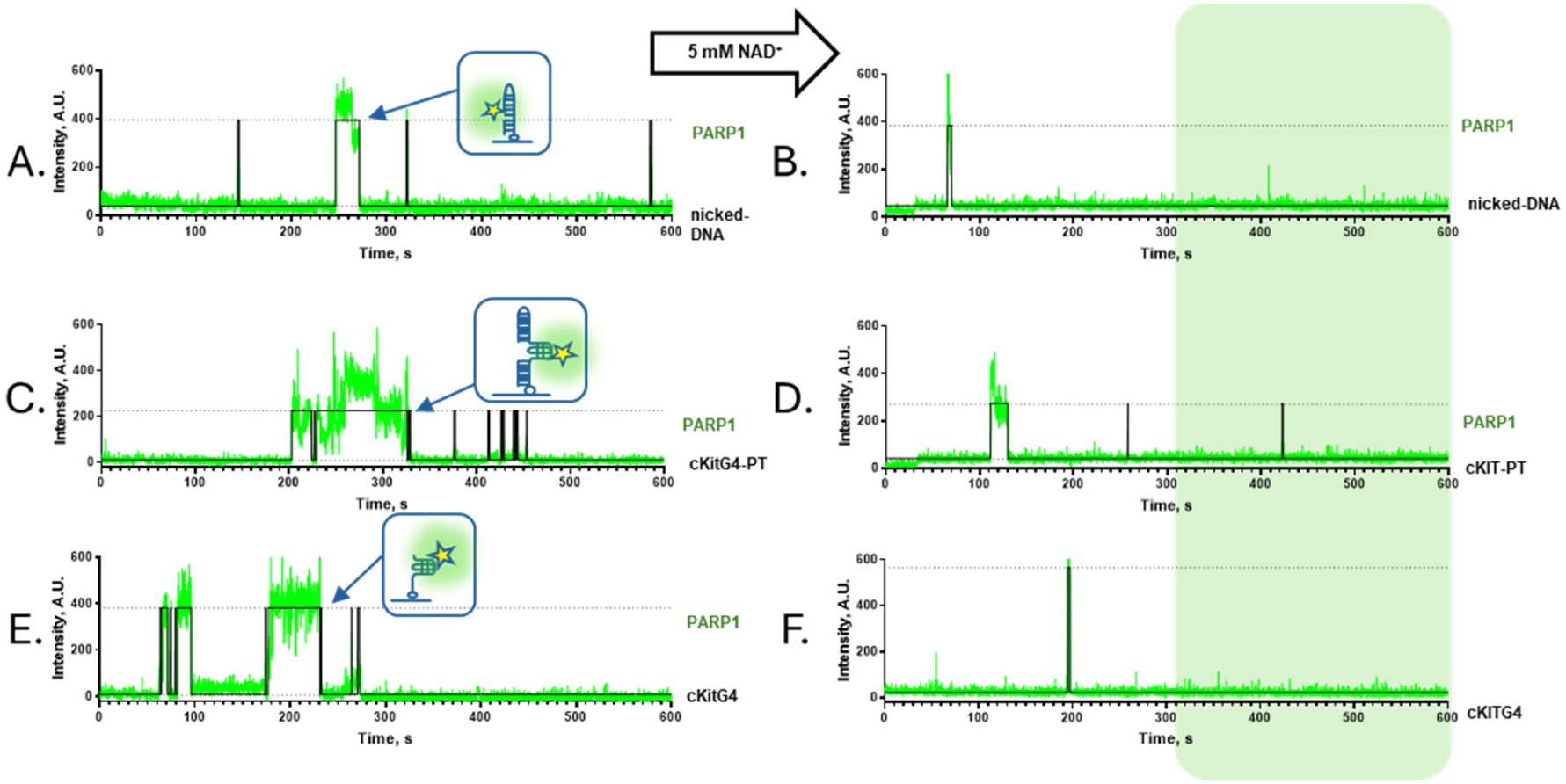
NAD^+^ Reduces PARP1 binding to DNA. Biotinylated partial duplex DNA containing the nicked **(A-B)**, cKITG4-PT **(C-D)** and cKITG4 **(E-F)** sequence were immobilized on a surface, while Cy3-labeled PARP1 (Cy3-PARP1) was infused into the reaction chamber (**A, C, and E**). In the second half panel, 5 mM NAD^+^ was infused in reaction chamber with Cy3-PARP1 (**B, D, and F)** which showed significantly a smaller number of binding events in the second half of the trajectory (Green shaded area in B, D and F).

## DISCUSSION

Our MP and smTIRFM experiments showed that PARP1 dynamically interacts with G4s and undergoes robust PARylation, especially when the substrates contain both G4 and a primer-template junction (represented by the cKITG4-PT). With MP experiments we show that PARP1 forms a 1:1 complex with both nicked and G4-PT DNA, and this 1:1 complex is sufficient for PARylation to occur. The 1:1 PARP1-DNA complex was verified using the 2-color smTIRFM experiments, where we saw that two color (Cy3 and Cy5) PARP1 molecules cannot coexist with the same DNA molecule and to bind on it they must compete.

Notably, in the presence of both nicked and cKITG4-PT DNA, accumulation of the PAR chains proceeded for several minutes, while the smTIRFM experiments suggested the “on” dwell times for the PARP1-DNA complexes of several seconds, which were further reduced in the presence of NAD^+^. Combined, these data suggest that the initial PARylation of the DNA-bound PARP1 results in the protein release from the DNA substrate and continuous enzymatic activity of the unbound protein. We noticed a more pronounced PARylation in the case of cKITG4-PT DNA compared to a canonical PARP1 substrate, namely nicked DNA, with up to ∼815 ADP-ribose units attached. The growth of PAR chains in the MP experiments continued for several minutes, which is much longer than the dwell times of the bound states recorded in the smTIRFM experiments. The most likely explanation is that the PARylation is activated by the PARP1 binding to the DNA substrate but continues after PARP1 dissociation. The change in the dwell times of cKITG4-PARP1 complexes observed in the smTIRFM experiments suggest that the simple cKITG4 can also promote PARylation, but the resulting chains were too short to be detected in the MP experiments. A recent study investigated the structure-specific functions of PAR by examining the effects of different branching lengths (54).

We can relate our findings to this study: we observed shorter PAR chains with the nicked DNA substrate, whereas the cKITG4-PT DNA substrate resulted in the formation of extensive PAR chains, suggesting that these structures may play diverse roles in cellular processes.

Single-molecule studies were conducted to examine the effects of PARPi on PARP1 using both nicked and G4-containing DNA substrates. Three potent PARPi were selected, inspired by a previously published study that categorized these inhibitors based on their allosteric effects on PARP1 (13). This classification defines PARPi into three types: Type I (proretention), which strongly enhance retention; Type II (modest proretention or no effect); and Type III (prorelease), which promote dissociation. Consistent with this framework, the Type I inhibitor EB-47 demonstrated enhanced retention of PARP1 on nicked DNA. Clinically relevant PARPi, such as Olaparib, were classified as Type II, exhibiting moderate retention, while Veliparib, a Type III inhibitor, was shown to promote dissociation (13). Our smTIRFM experiments using biotinylated nicked DNA substrates revealed similar trends. EB-47 and Olaparib increased PARP1 retention times, while Veliparib facilitated increased dissociation, aligning with their respective classifications. Additionally, we observed a pattern of tighter binding affinities (Kds) and longer dwell times for PARP1 when interacting with nicked DNA substrates, which mimic single-strand breaks. EB-47 showed the strongest trapping activity, followed by Olaparib and Veliparib, indicating varying degrees of retention effects (see **Table 1**). These findings are consistent with a study that used fluorescence polarization assays to calculate equilibrium dissociation constants (Kds) of PARP1 binding to SSB DNA with and without PARPi. Without an inhibitor, the Kd was ∼90 nM, while Veliparib increased it to ∼300 nM (∼3.4× difference), Olaparib reduced it to ∼60 nM (∼0.67× difference), and EB-47 drastically reduced it to ∼5 nM (∼0.06× difference) (13). This trend closely aligns with our calculated Kds values (see Table 1), where EB-47 significantly enhances PARP1 binding/trapping, followed by Olaparib and Veliparib promoting release. Furthermore, dissociation rate constants (kd) of PARP1 from nicked DNA substrate were measured using surface plasmon resonance in the presence or absence of the PARPi, for EB-47, Olaparib, and Veliparib, the kd values changed by ∼0.34×, ∼0.69×, and ∼1.42×, respectively, showing a similar trend to our data (EB-47 > Olaparib > Veliparib, see Table 1) (13). This consistency highlights the distinct allosteric effects of each PARPi on PARP1 trapping and dissociation dynamics.

A completely different effect was observed with the G4-containing DNAs. Our analyses utilized two configurations of the G4 DNA substrate: the cKITG4, which was a small DNA substrate and only contained a G-quadruplex, and the cKITG4-PT, which was larger and consisting of G4 and two dsDNA arms one of which represented a primer-template junction. While initially, the cKITG4-PT was designed to simply increase the substrate size for the MP experiments, it revealed different complexation and ability to activate PARylation compared to its smaller counterpart. In cells, the replication-stalling G4s are expected to have features similar to our cKITG4-PT and therefore to strongly activate PARP1. In the smTIRFM experiments, which are not limited by the size of DNA biomolecules, we observed a ∼80x tighter Kd for cKITG4 DNA substrate compared to cKITG4-PT. In case of cKITG4 DNA with PARPi, we observed ∼4x, ∼7x and ∼30x higher Kds (Veliparib, Olaparib and EB-47) compared to no PARPi. Notably, this reduction in affinity was due to the slower association rates and not to the change in the stability of the cKITG4-PARP1 complex. Differently from both cKITG4 and nicked DNA, none of the three inhibitors affected PARP1 binding to the cKITG4-PT DNA.

PARPi inhibitors exhibit pleiotropic activities. PARPi can either bind to catalytic site on PARP1, preventing NAD^+^ binding and PARylation or allosteric trapping on DNA substrate in cells, however different inhibitors differ in their capacity (9,13,42,55,56). Recognizing effects of PARPi on different PARP1-G4 complexed vs. other lesions recognized by PARP1 could potentially lead to the development of more effective anti-cancer drugs (55).

It was shown recently that FANCJ helicase may help to activate PARP1 (57). In FANCJ deficient cells, PARP1 becomes trapped on G4 DNA, reducing cell sensitivity to PARPi. The same study also showed that interaction between FANCJ and MMR proteins is crucial for PARP1 activity. Without MSH2, cells become more sensitive to PARPi. In BRCA1-deficient cells, losing FANCJ is like losing or inhibiting PARP1. This emphasizes the importance of PARP1 activity during DNA replication for these cells. It was proposed that the effectiveness of PARPi in BRCA1-deficient cancers may be due to inhibiting PARP1 activity during s-phase of cell division rather than trapping it on DNA which was thought to be the mechanism earlier. Notably, FANCJ has a capacity to recognize the replication stalling G4s (58), and to both unfold and refold them (59). The FANCJ activity at these G4s may help to maintain their presence until they are either replicated through or recruit PARP1 and trigger PARylation signaling. On the other hand, PARP1 interaction with G4s may act as a signal to G4 other specific helicases including BLM, WRN, and DNA2, which all have specificity for different DNA G4 structures (60). This facilitates rapid sampling of the G4s to bring the best helicase to process the G4s. PARP1 dissociates from the G4s and a helicase, in conjunction with a non-replicative polymerase resolves the replication block.

## SUPPLEMENTARY DATA

Supplementary Data are available at NAR online.

## AUTHOR CONTRIBUTIONS

PG: Expression, purification, preparation of fluorescently labeled PARP1, smTIRFM, MP based experiments, circular dichroism, data analysis, writing manuscript and conceptual design of the study. F.E.B: Expression, purification, preparation of fluorescently labeled PARP1, smTIRFM experiments and conceptual design of the study. RM: MP based experiments. MS: Conceptual design, data analysis, interpretation, writing manuscript, supervision and funding acquisition.

## ACKNOWLEDGEMENTS

We thank Dr. Kevin D. Raney from the Department of Biochemistry and Molecular Biology, University of Arkansas for Medical Sciences for generously providing the PARP1 expression construct.

## REFERENCES

1. Durkacz, B.W., Omidiji, O., Gray, D.A. and Shall, S. (1980) (ADP-ribose)n participates in DNA excision repair. Nature, 283, 593–596.

2. Hayaishi, O. and Ueda, K. (1977) Poly(ADP-ribose) and ADP-ribosylation of proteins. Annu Rev Biochem, 46, 95–116.

3. Hilz, H. and Stone, P. (1976) Poly(ADP-ribose) and ADP-ribosylation of proteins. Rev Physiol Biochem Pharmacol, 76, 1–58, 177.

4. Alemasova, E.E. and Lavrik, O.I. (2019) Poly(ADP-ribosyl)ation by PARP1: reaction mechanism and regulatory proteins. Nucleic Acids Res, 47, 3811–3827.

5. Martin-Hernandez, K., Rodriguez-Vargas, J.M., Schreiber, V. and Dantzer, F. (2017) Expanding functions of ADP-ribosylation in the maintenance of genome integrity. Semin Cell Dev Biol, 63, 92–101.

6. Ray Chaudhuri, A. and Nussenzweig, A. (2017) The multifaceted roles of PARP1 in DNA repair and chromatin remodelling. Nat Rev Mol Cell Biol, 18, 610–621.

7. Spiegel, J.O., Van Houten, B. and Durrant, J.D. (2021) PARP1: Structural insights and pharmacological targets for inhibition. DNA Repair (Amst), 103, 103125.

8. Langelier, M.F. and Pascal, J.M. (2013) PARP-1 mechanism for coupling DNA damage detection to poly(ADP-ribose) synthesis. Curr Opin Struct Biol, 23, 134–143.

9. Satoh, M.S. and Lindahl, T. (1992) Role of poly(ADP-ribose) formation in DNA repair. Nature, 356, 356–358.

10. Eustermann, S., Wu, W.F., Langelier, M.F., Yang, J.C., Easton, L.E., Riccio, A.A., Pascal, J.M. and Neuhaus, D. (2015) Structural Basis of Detection and Signaling of DNA Single-Strand Breaks by Human PARP-1. Mol Cell, 60, 742–754.

11. Rouleau-Turcotte, E., Krastev, D.B., Pettitt, S.J., Lord, C.J. and Pascal, J.M. (2022) Captured snapshots of PARP1 in the active state reveal the mechanics of PARP1 allostery. Mol Cell, 82, 2939–2951 e2935.

12. Barkauskaite, E., Jankevicius, G. and Ahel, I. (2015) Structures and Mechanisms of Enzymes Employed in the Synthesis and Degradation of PARP-Dependent Protein ADP-Ribosylation. Mol Cell, 58, 935–946.

13. Zandarashvili, L., Langelier, M.F., Velagapudi, U.K., Hancock, M.A., Steffen, J.D., Billur, R., Hannan, Z.M., Wicks, A.J., Krastev, D.B., Pettitt, S.J. et al. (2020) Structural basis for allosteric PARP-1 retention on DNA breaks. Science, 368.

14. Hottiger, M.O., Hassa, P.O., Luscher, B., Schuler, H. and Koch-Nolte, F. (2010) Toward a unified nomenclature for mammalian ADP-ribosyltransferases. Trends Biochem Sci, 35, 208–219.

15. Tao, Z., Gao, P. and Liu, H.W. (2009) Identification of the ADP-ribosylation sites in the PARP-1 automodification domain: analysis and implications. J Am Chem Soc, 131, 14258–14260.

16. Gagne, J.P., Ethier, C., Defoy, D., Bourassa, S., Langelier, M.F., Riccio, A.A., Pascal, J.M., Moon, K.M., Foster, L.J., Ning, Z. et al. (2015) Quantitative site-specific ADP-ribosylation profiling of DNA-dependent PARPs. DNA Repair (Amst), 30, 68–79.

17. Gellert, M., Lipsett, M.N. and Davies, D.R. (1962) Helix formation by guanylic acid. Proceedings of the National Academy of Sciences of the United States of America, 48, 2013–2018.

18. Burge, S., Parkinson, G.N., Hazel, P., Todd, A.K. and Neidle, S. (2006) Quadruplex DNA: sequence, topology and structure. Nucleic Acids Res, 34, 5402–5415.

19. Víglaský, V., Bauer, L. and Tlučková, K. (2010) Structural Features of Intra- and Intermolecular G-Quadruplexes Derived from Telomeric Repeats. Biochemistry, 49, 2110–2120.

20. Lane, A.N., Chaires, J.B., Gray, R.D. and Trent, J.O. (2008) Stability and kinetics of G-quadruplex structures. Nucleic acids research, 36, 5482–5515.

21. Edwards, A.D., Marecki, J.C., Byrd, A.K., Gao, J. and Raney, K.D. (2021) G-Quadruplex loops regulate PARP-1 enzymatic activation. Nucleic Acids Res, 49, 416–431.

22. Gray, R.D. and Chaires, J.B. (2008) Kinetics and mechanism of K+- and Na+-induced folding of models of human telomeric DNA into G-quadruplex structures. Nucleic acids research, 36, 4191–4203.

23. Li, X., Wang, J., Gong, X., Zhang, M., Kang, S., Shu, B., Wei, Z., Huang, Z.S. and Li, D. (2020) Upregulation of BCL-2 by acridone derivative through gene promoter i-motif for alleviating liver damage of NAFLD/NASH. Nucleic Acids Res, 48, 8255–8268.

24. You, J., Li, H., Lu, X.M., Li, W., Wang, P.Y., Dou, S.X. and Xi, X.G. (2017) Effects of monovalent cations on folding kinetics of G-quadruplexes. Biosci Rep, 37.

25. Bhattacharyya, D., Mirihana Arachchilage, G. and Basu, S. (2016) Metal Cations in G-Quadruplex Folding and Stability. Front Chem, 4, 38.

26. Ma, Y., Iida, K. and Nagasawa, K. (2020) Topologies of G-quadruplex: Biological functions and regulation by ligands. Biochem Biophys Res Commun, 531, 3–17.

27. Iyer, D.R. and Rhind, N. (2017) Replication fork slowing and stalling are distinct, checkpoint-independent consequences of replicating damaged DNA. PLOS Genetics, 13, e1006958.

28. Sato, K., Martin-Pintado, N., Post, H., Altelaar, M. and Knipscheer, P. (2021) Multistep mechanism of G-quadruplex resolution during DNA replication. Sci Adv, 7, eabf8653.

29. Sato, K. and Knipscheer, P. (2023) G-quadruplex resolution: From molecular mechanisms to physiological relevance. DNA Repair (Amst), 130, 103552.

30. Soldatenkov, V.A., Vetcher, A.A., Duka, T. and Ladame, S. (2008) First evidence of a functional interaction between DNA quadruplexes and poly(ADP-ribose) polymerase-1. ACS Chem Biol, 3, 214–219.

31. Hanuman Singh, D., Deeksha, W. and Rajakumara, E. (2024) Characterization of PARP1 binding to c-KIT1 G-quadruplex DNA: Insights into domain-specific interactions. Biophys Chem, 315, 107330.

32. Soldatenkov, V.A., Vetcher, A.A., Duka, T. and Ladame, S. (2008) First Evidence of a Functional Interaction Between DNA Quadruplexes and Poly(ADP-Ribose) Polymerase-1. Acs Chemical Biology.

33. Langelier, M.F., Planck, J.L., Servent, K.M. and Pascal, J.M. (2011) Purification of human PARP-1 and PARP-1 domains from Escherichia coli for structural and biochemical analysis. Methods Mol Biol, 780, 209–226.

34. Gaur, P., Bain, F.E., Honda, M., Granger, S.L. and Spies, M. (2023) Single-Molecule Analysis of the Improved Variants of the G-Quadruplex Recognition Protein G4P. Int J Mol Sci, 24.

35. Fairlamb, M.S., Whitaker, A.M., Bain, F.E., Spies, M. and Freudenthal, B.D. (2021) Construction of a Three-Color Prism-Based TIRF Microscope to Study the Interactions and Dynamics of Macromolecules. Biology (Basel), 10.

36. Bain, F.E., Fischer, L.A., Chen, R. and Wold, M.S. (2018) Single-Molecule Analysis of Replication Protein A-DNA Interactions. Methods Enzymol, 600, 439–461.

37. Hon, J. and Gonzalez, R.L., Jr. (2019) Bayesian-Estimated Hierarchical HMMs Enable Robust Analysis of Single-Molecule Kinetic Heterogeneity. Biophys J, 116, 1790–1802.

38. Tibbs, J., Ghoneim, M., Caldwell, C.C., Buzynski, T., Bowie, W., Boehm, E.M., Washington, M.T., Tabei, S.M.A. and Spies, M. (2021) KERA: analysis tool for multi-process, multi-state single-molecule data. Nucleic Acids Res, 49, e53.

39. Fernando, H., Reszka, A.P., Huppert, J., Ladame, S., Rankin, S., Venkitaraman, A.R., Neidle, S. and Balasubramanian, S. (2006) A conserved quadruplex motif located in a transcription activation site of the human c-kit oncogene. Biochemistry, 45, 7854–7860.

40. Zheng, K.W., Zhang, J.Y., He, Y.D., Gong, J.Y., Wen, C.J., Chen, J.N., Hao, Y.H., Zhao, Y. and Tan, Z. (2020) Detection of genomic G-quadruplexes in living cells using a small artificial protein. Nucleic Acids Res, 48, 11706–11720.

41. Ivanov, V.I., Minchenkova, L.E., Schyolkina, A.K. and Poletayev, A.I. (1973) Different Conformations of Double-Stranded Nucleic-Acid in Solution as Revealed by Circular-Dichroism. Biopolymers, 12, 89–110.

42. Xue, H., Bhardwaj, A., Yin, Y., Fijen, C., Ephstein, A., Zhang, L., Ding, X., Pascal, J.M., VanArsdale, T.L. and Rothenberg, E. (2022) A two-step mechanism governing PARP1-DNA retention by PARP inhibitors. Sci Adv, 8, eabq0414.

43. Asor, R. and Kukura, P. (2022) Characterising biomolecular interactions and dynamics with mass photometry. Curr Opin Chem Biol, 68, 102132.

44. Young, G. and Kukura, P. (2019) Interferometric Scattering Microscopy. Annual review of physical chemistry, 70, 301–322.

45. Ashworth, A. and Lord, C.J. (2018) Synthetic lethal therapies for cancer: what’s next after PARP inhibitors? Nat Rev Clin Oncol, 15, 564–576.

46. Helleday, T. (2011) The underlying mechanism for the PARP and BRCA synthetic lethality: Clearing up the misunderstandings. Mol Oncol, 5, 387–393.

47. Tuli, R., Shiao, S.L., Nissen, N., Tighiouart, M., Kim, S., Osipov, A., Bryant, M., Ristow, L., Placencio-Hickok, V., Hoffman, D. et al. (2019) A phase 1 study of veliparib, a PARP-1/2 inhibitor, with gemcitabine and radiotherapy in locally advanced pancreatic cancer. EBioMedicine, 40, 375–381.

48. Jagtap, P.G., Southan, G.J., Baloglu, E., Ram, S., Mabley, J.G., Marton, A., Salzman, A. and Szabo, C. (2004) The discovery and synthesis of novel adenosine substituted 2,3-dihydro-1H-isoindol-1-ones: potent inhibitors of poly(ADP-ribose) polymerase-1 (PARP-1). Bioorg Med Chem Lett, 14, 81–85.

49. Kamaletdinova, T., Fanaei-Kahrani, Z. and Wang, Z.-Q. (2019) The Enigmatic Function of PARP1: From PARylation Activity to PAR Readers. Cells, 8, 1625.

50. Wei, H. and Yu, X. (2016) Functions of PARylation in DNA Damage Repair Pathways. *Genomics*, Proteomics & Bioinformatics, 14, 131–139.

51. Krüger, A., Bürkle, A., Hauser, K. and Mangerich, A. (2020) Real-time monitoring of PARP1-dependent PARylation by ATR-FTIR spectroscopy. Nature Communications, 11, 2174.

52. Soldatenkov, V.A., Vetcher, A.A., Duka, T. and Ladame, S. (2008) First Evidence of a Functional Interaction between DNA Quadruplexes and Poly(ADP-ribose) Polymerase-1. ACS Chemical Biology, 3, 214–219.

53. Lonskaya, I., Potaman, V.N., Shlyakhtenko, L.S., Oussatcheva, E.A., Lyubchenko, Y.L. and Soldatenkov, V.A. (2005) Regulation of poly(ADP-ribose) polymerase-1 by DNA structure-specific binding. J Biol Chem, 280, 17076–17083.

54. Aberle, L., Kruger, A., Reber, J.M., Lippmann, M., Hufnagel, M., Schmalz, M., Trussina, I., Schlesiger, S., Zubel, T., Schutz, K. et al. (2020) PARP1 catalytic variants reveal branching and chain length-specific functions of poly(ADP-ribose) in cellular physiology and stress response. Nucleic Acids Res, 48, 10015–10033.

55. Macgilvary, N. and Cantor, S.B. (2024) Positioning loss of PARP1 activity as the central toxic event in BRCA-deficient cancer. DNA Repair, 144.

56. Rudolph, J., Jung, K. and Luger, K. (2022) Inhibitors of PARP: Number crunching and structure gazing. Proc Natl Acad Sci U S A, 119, e2121979119.

57. Cong, K., MacGilvary, N., Lee, S., Macleod, S.G., Calvo, J., Peng, M., Kousholt, A., Day, T.A. and Cantor, S.B. (2024) FANCJ promotes PARP1 activity during DNA replication that is essential in deficient cells. Nature Communications, 15.

58. Sarkies, P., Murat, P., Phillips, L.G., Patel, K.J., Balasubramanian, S. and Sale, J.E. (2011) FANCJ coordinates two pathways that maintain epigenetic stability at G-quadruplex DNA. Nucleic Acids Res, 40, 1485–1498.

59. Wu, C.G. and Spies, M. (2016) G-quadruplex recognition and remodeling by the FANCJ helicase. Nucleic Acids Res, 44, 8742–8753.

60. Ray, S., Tillo, D., Boer, R.E., Assad, N., Barshai, M., Wu, G., Orenstein, Y., Yang, D., Schneekloth, J.S. and Vinson, C. (2020) Custom DNA Microarrays Reveal Diverse Binding Preferences of Proteins and Small Molecules to Thousands of G-Quadruplexes. ACS Chemical Biology, 15, 925–935.

