## Supplementary Figures and Tables for "Single-molecule analysis of PARP1-G-quadruplex interaction"

**Supplementary Table 1. Oligonucleotides used in this study**

| Name | DNA sequence | size | technique |
|---|---|---|---|
| Nicked | 5'-CATGTAGTCACTATGAGGCATAGTGACTACATGA TCTAATGCATTAAGTTCCTTAATGCATTAGAT-3' | 66 nt | MP |
| cKITG4-PT | 5'- CATGTAGTCACTATGATTAGGCATAGTGACTACATGGGG CGGGCGCGAGGGAGGGGAGGATCTAATGCATTAAGTTCGCT TAATGCATTAGAT -3' | 93 nt | MP |
| cKITG4 | 5'-**GGG**C**GGG**CGCGA**GGG**A**GGGG**AGG | 23 nt | MP |
| ssDNA | 5'-(dT)$_{90}$-3' | 90 nt | MP |
| cKITG4-PT | 5'- TGGCGACGGCAGCGAGGCGGGCGGGCGCGAGGGAGG GGAGGTCTAATGCATTAAGTTCCTTAATGCATTAGA -3' | 72 nt | smTIRFM |
| nicked | 5'- TGGCGACGGCAGCGAGGCTCTAATGCATTAAGTTCCTT AATGCATTAGA -3' | 49 nt | smTIRFM |
| Biotin-base | 5'- GCCTCGCTGCCGTCGCCA/3Bio/-3' | 18 nt | smTIRFM |
| dsDNA | 5'-CATGTAGTCACTATGCTACTGTG/3Bio/-3' 3'-GTACATCAGTGATACGATGACAC-5' | 23 nt | smTIRFM |
| cMYCG4 | 5'-TGGGTGGGTAGGGTGGGTTT/3Bio/-3' | 20 nt | smTIRFM |
| hTelG4 | 5'-TTAGGGTTAGGGTTAGGGTTAGGG/3Bio/-3' | 24 nt | smTIRFM |
| cKITG4 | 5'- GGGCGGGCGCGAGGGAGGGGAGG/3Bio/-3' | 23 nt | smTIRFM |
| ssDNA | 5'- CATGTAGTCACTATGCTACTGTG/3Bio/-3' | 23 nt | smTIRFM |

The biotin-base oligo and its complementary sequence on the smTIRFM nicked and cKITG4-PT substrates are highlighted in purple. The cKIT G-quadruplex forming sequence is highlighted in orange.

`

A.

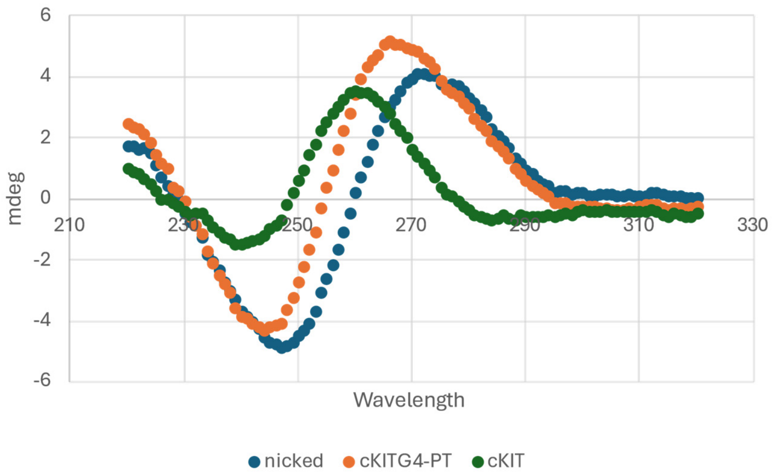


B.

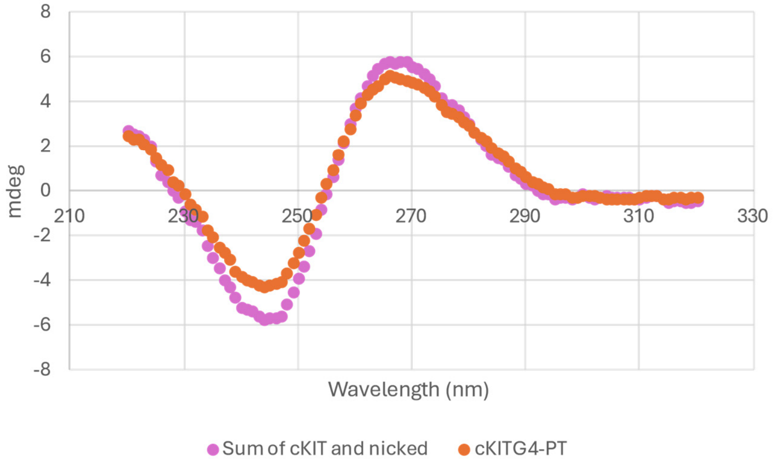


**Supplementary Figure 1.** Circular Dichroism (CD) spectra for folded DNA substrates: nicked, cKITG4-PT, and cKIT (see Supplementary Table 1). Samples were prepared in buffer containing 20 mM Tris, 100 mM KCl, 10 mM $MgCl_2$, and 1 mM EDTA at a concentration of 5 µM. DNA samples were heat-denatured at 95°C for 5 minutes and then slowly cooled to allow secondary structure formation. CD measurements were conducted at room temperature using a JASCO J-810 spectropolarimeter over the wavelength range of 220–320 nm, with an average of 10 scans per sample. (**A**) CD spectra of cKIT, nicked, and cKITG4-PT substrates are shown. (**B)** Comparison of the CD spectrum of cKITG4-PT with the sum of cKIT and nicked spectra, demonstrating similarity to the cKIT-PT spectrum.

`

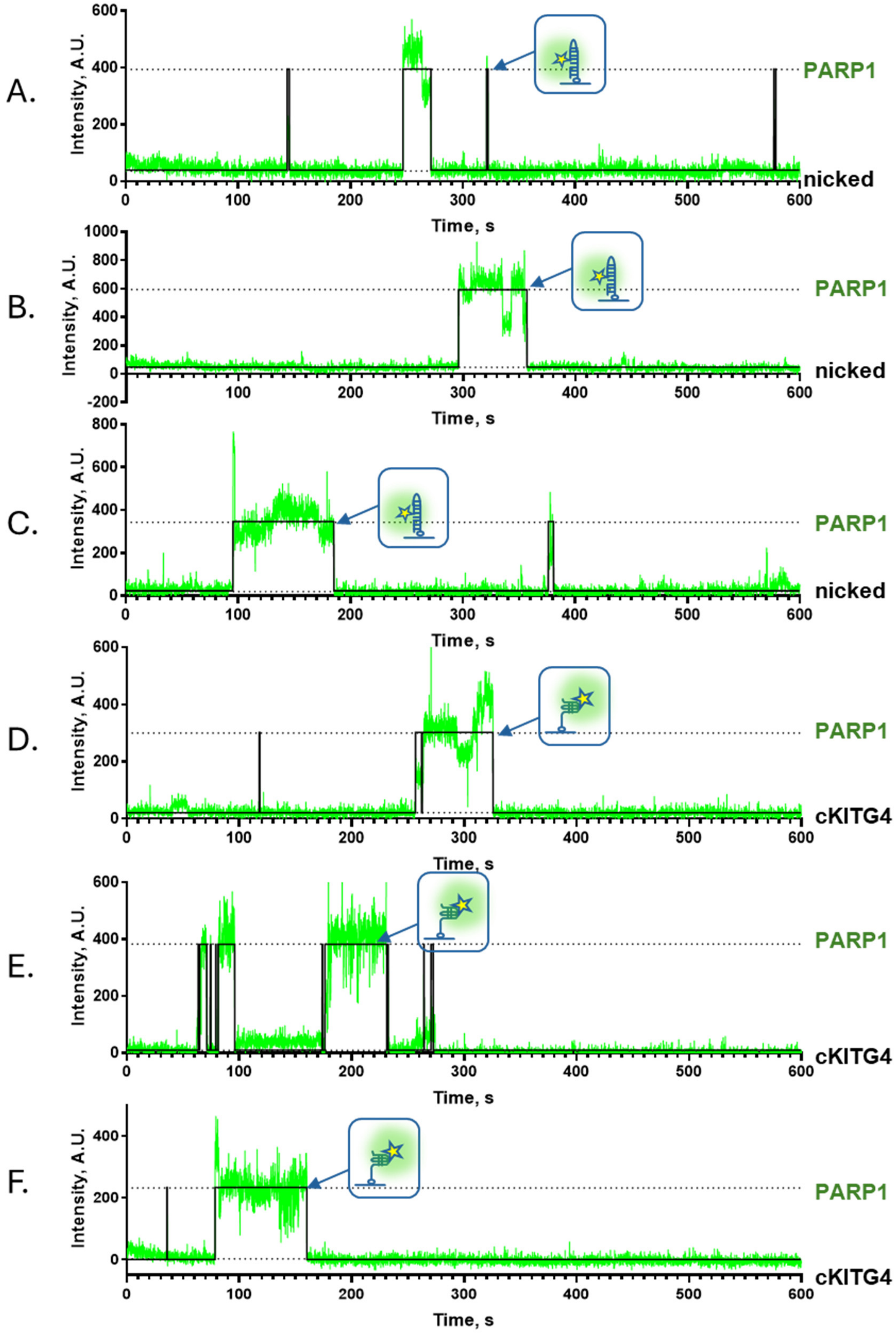


**Supplementary Figure 2.** Representative smTIRFM trajectories of Cy3-labeled PARP1 binding to surface-tethered nicked DNA (A-C) and cKITG4 DNA (D-F). Biotinylated DNA constructs (100 pM) were immobilized on the surface, while Cy3-labeled PARP1 was infused at 100 pM for the nicked DNA substrate and 200 pM for the cKITG4 DNA substrate. Representative fluorescence trajectories (green) are shown with their corresponding idealized fits (black), illustrating the binding interactions.

`

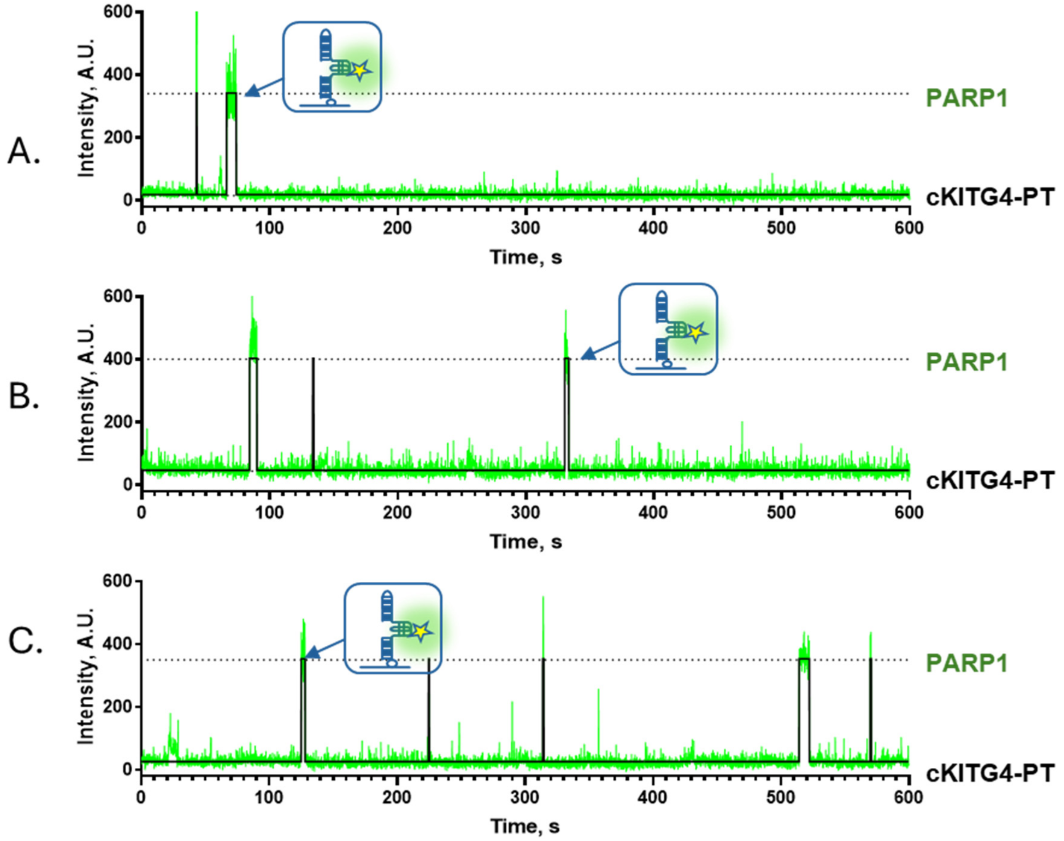


**Supplementary Figure 3.** Representative smTIRFM trajectories of Cy3-labeled PARP1 binding to surface-tethered cKITG4-PT (A-C). Biotinylated DNA constructs (100 pM) were immobilized on a surface, while 100 pM Cy3-PARP1 were infused into the reaction chamber. The representative trajectories show a fluorescence trajectory (green) overlaid with an idealized trajectory (black).

`

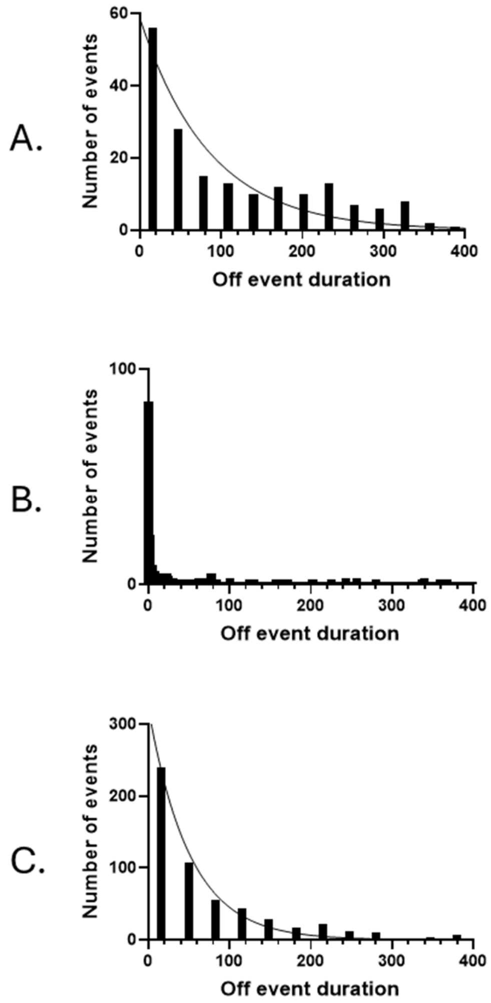


**Supplementary Figure 4.** Dwell-time distributions (smTIRFM experiment) for nicked, cKITG4 and cKITG4-PT DNA substrates which were constructed from all "ON" dwell times and fitted using a single-exponential function.

`

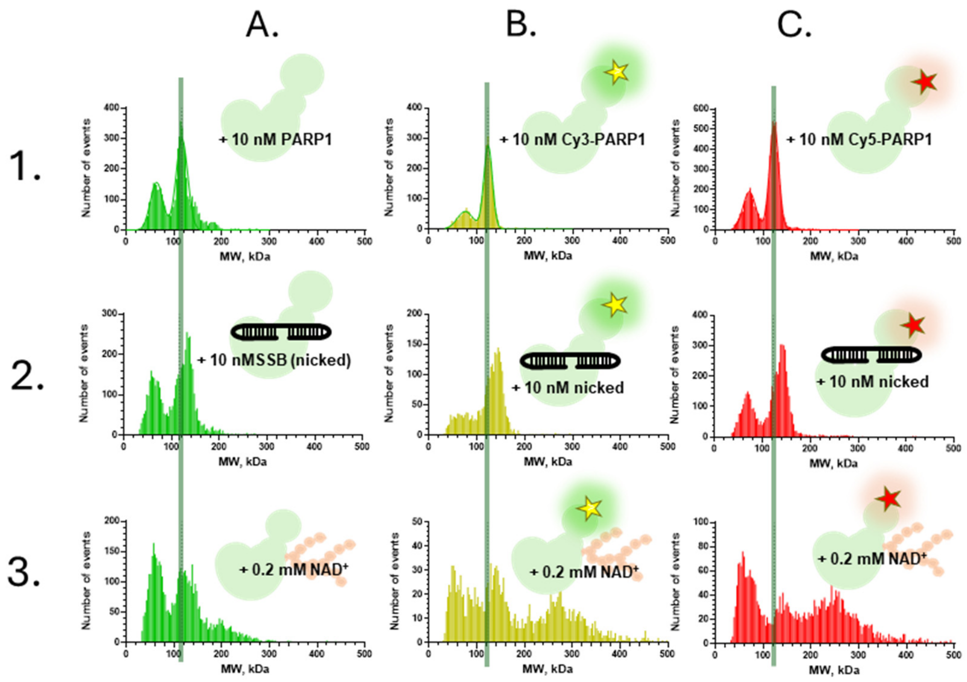


**Supplementary Figure 5.** Mass photometry analysis of DNA substrate interactions with PARP1 in presence of PARPi. Panels A-B and C-D depicts nicked and cKIT-PT DNA substrates, revealing a molecular weight shift (indicated by the transition from a gray line) upon PARP1 addition. This shift suggests the formation of PARP1-nicked and PARP1-cKIT-PT complexes. Following the introduction of 0.2 mM $NAD^+$, there is no formation of high molecular weight PAR chains, demonstrating the PARylation process is inhibited in each case A-B (nicked) and C-D (cKITG4-PT).

`

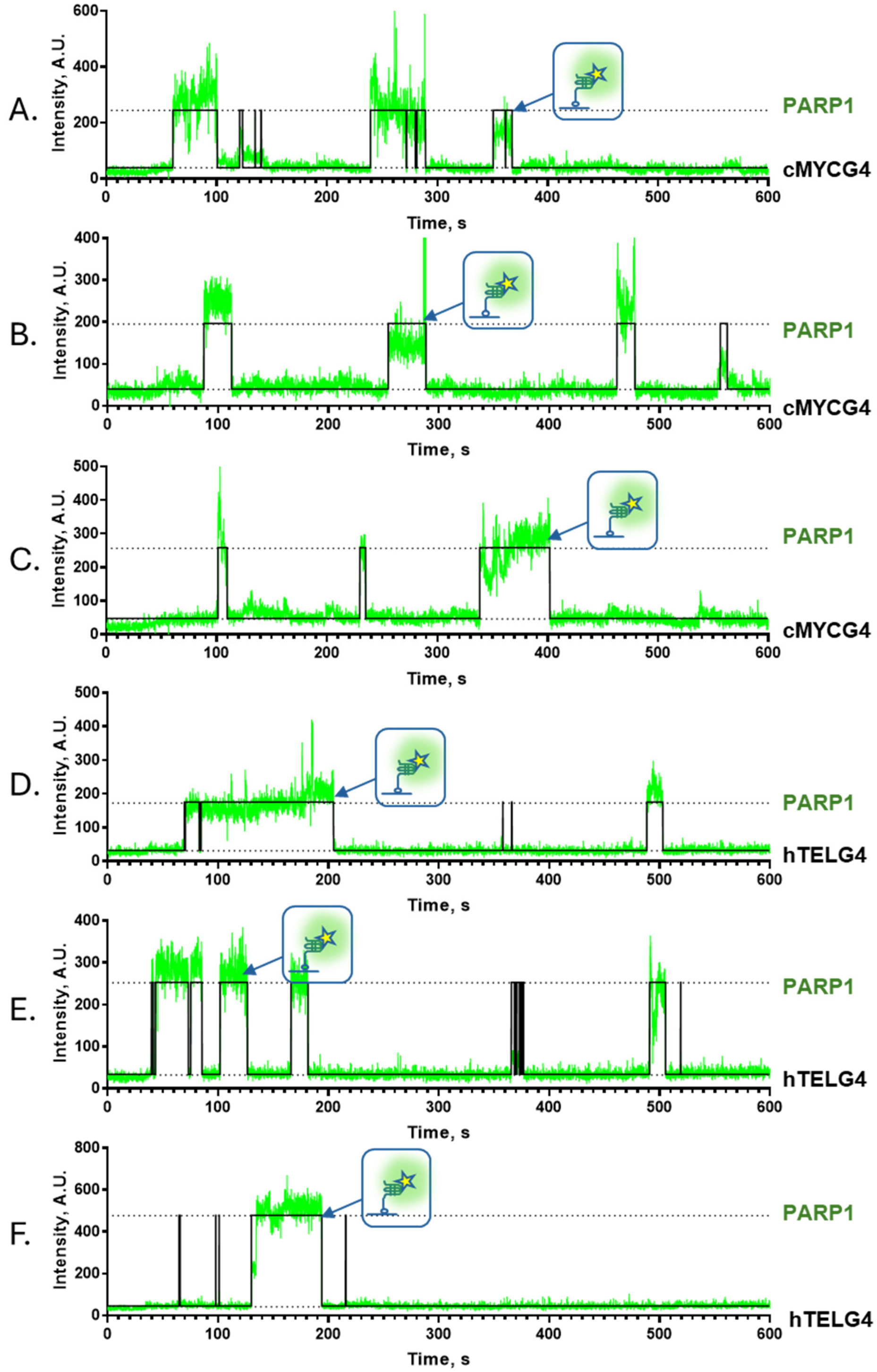


**Supplementary Figure 6.** Representative smTIRFM trajectories of Cy3-labeled PARP1 binding to surface-tethered cMYC G4 (A-C) and hTEL G4 (D-F) DNA. Biotinylated DNA constructs (100 pM) were immobilized on the surface, while Cy3-PARP1 was infused into the reaction chamber at 200 pM for the cMYC G4 substrate and 500 pM for the hTEL G4 substrate. Representative trajectories display fluorescence intensity (green) overlaid with idealized fits (black), illustrating binding dynamics.

`

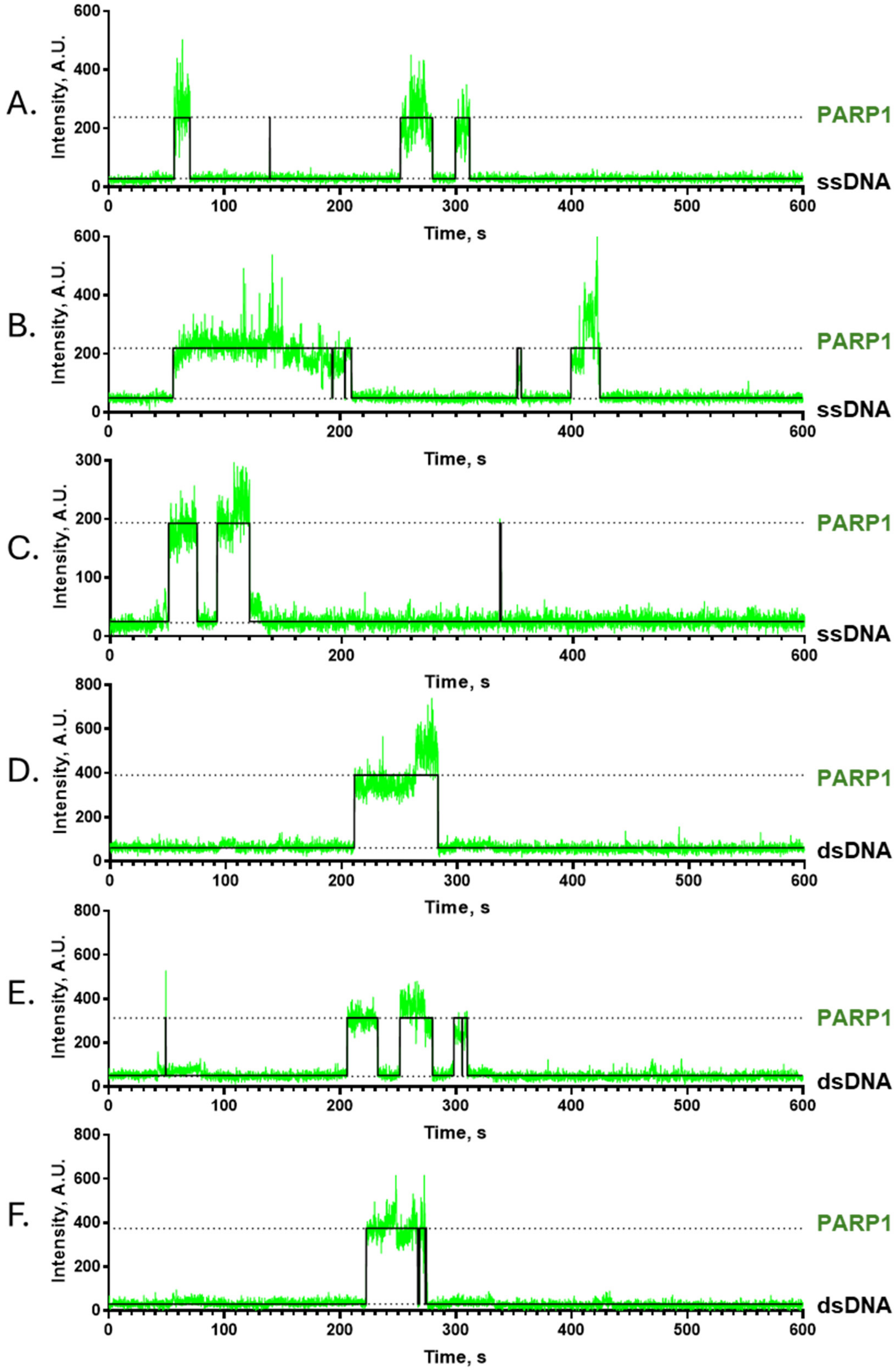


**Supplementary Figure 7.** Representative smTIRFM trajectories of Cy3-labeled PARP1 binding to surface-tethered ssDNA (A-C) and dsDNA (D-F) DNA. Biotinylated DNA constructs (100 pM) were immobilized on a surface, while 500 pM of Cy3-PARP1 were infused into the reaction chamber with ssDNA and 200 pM in chamber with dsDNA. The representative trajectories show a fluorescence trajectory (green) overlaid with an idealized trajectory (black).

`

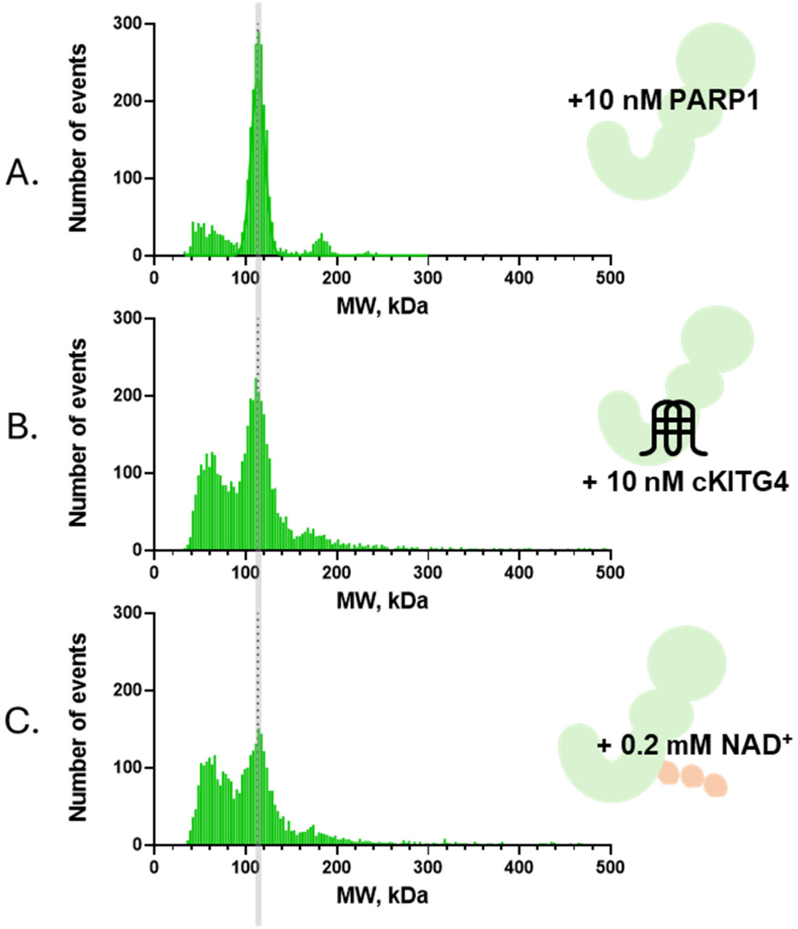


**Supplementary Figure 8**. Mass Photometry analysis of the interaction between cKITG4 DNA and PARP1. (A) Mass Photometry measurement of PARP1 alone (B) Addition of cKIT G4 DNA did not result in a detectable shift in molecular weight due to the relatively small size of the cKIT DNA substrate. (C) Subsequent addition of 0.2 mM $NAD^+$ initiated PARylation, providing further insight into the interaction dynamics.

`

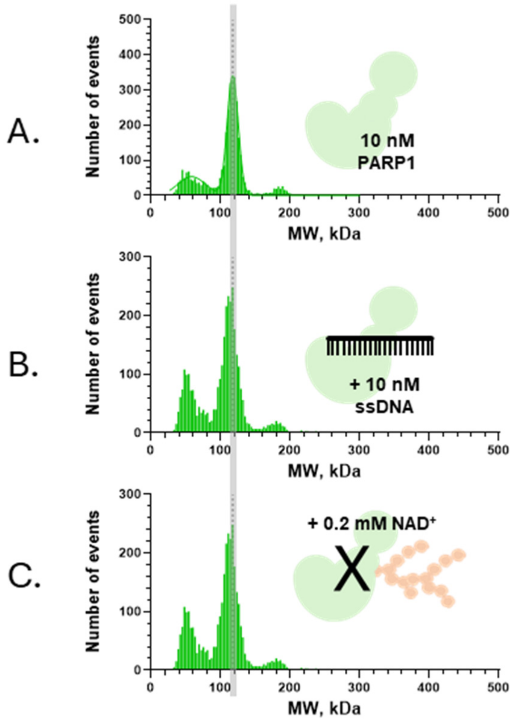


**Supplementary Figure 9**. Mass Photometry analysis of the interaction between single stranded DNA and PARP1. (A) Mass Photometry measurement of PARP1 alone (B) Addition of ssDNA did not result in a detectable shift in molecular weight. (C) Subsequent addition of 0.2 mM $NAD^+$ did not initiate PARylation.

`

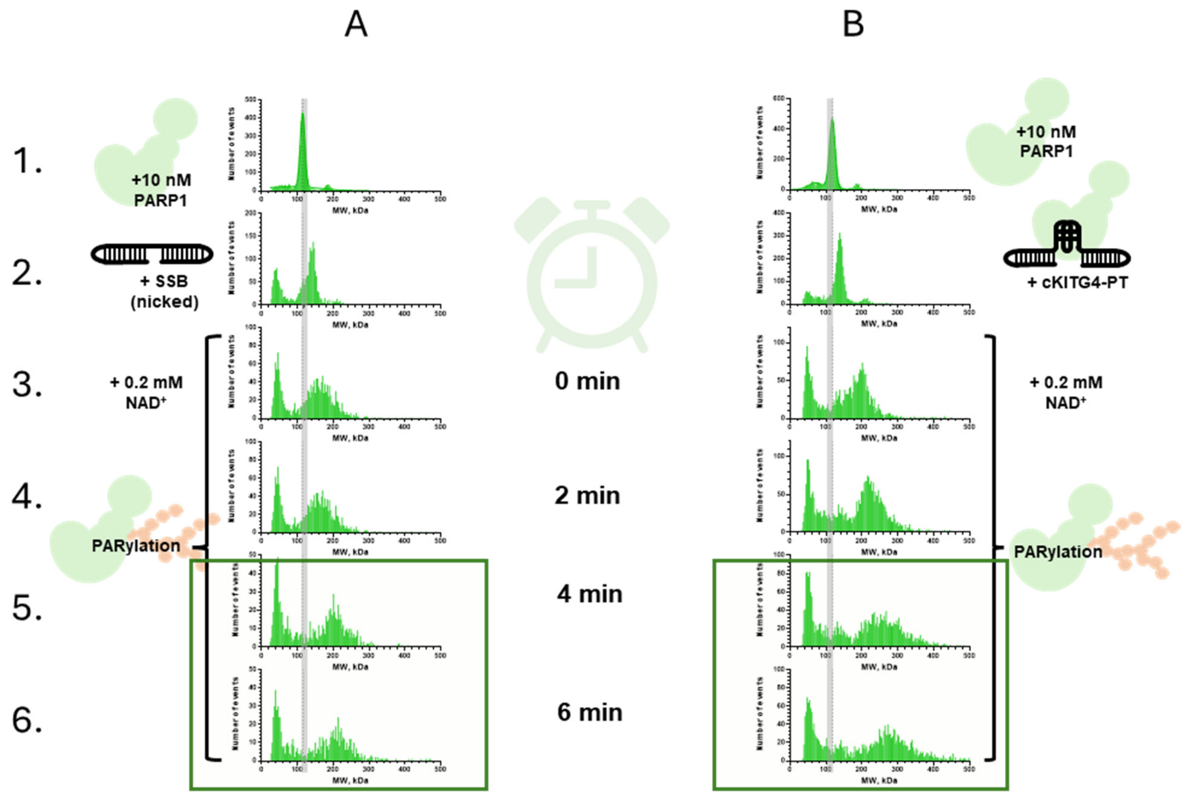


**Supplementary Figure 10.** Kinetics of PARylation. Panel A illustrates PARP1-mediated PARylation in the presence of a nicked DNA substrate, while Panel B demonstrates PARylation with the cKITG4-PT DNA substrate. Subpanels A1 and B1 depict 10 nM PARP1 alone. In subpanel A2, 10 nM nicked DNA substrate is added, and in subpanel B2, 10 nM cKITG4-PT DNA substrate is introduced. Subpanels A3 and B3 show the initiation of PARylation upon the addition of 0.2 mM $NAD^+$. The progression of PARylation over time is demonstrated in subpanels A4 and B4, showing intermediate PAR chain formation, which reaches saturation in subpanels A5–A6 and B5–B6.

`

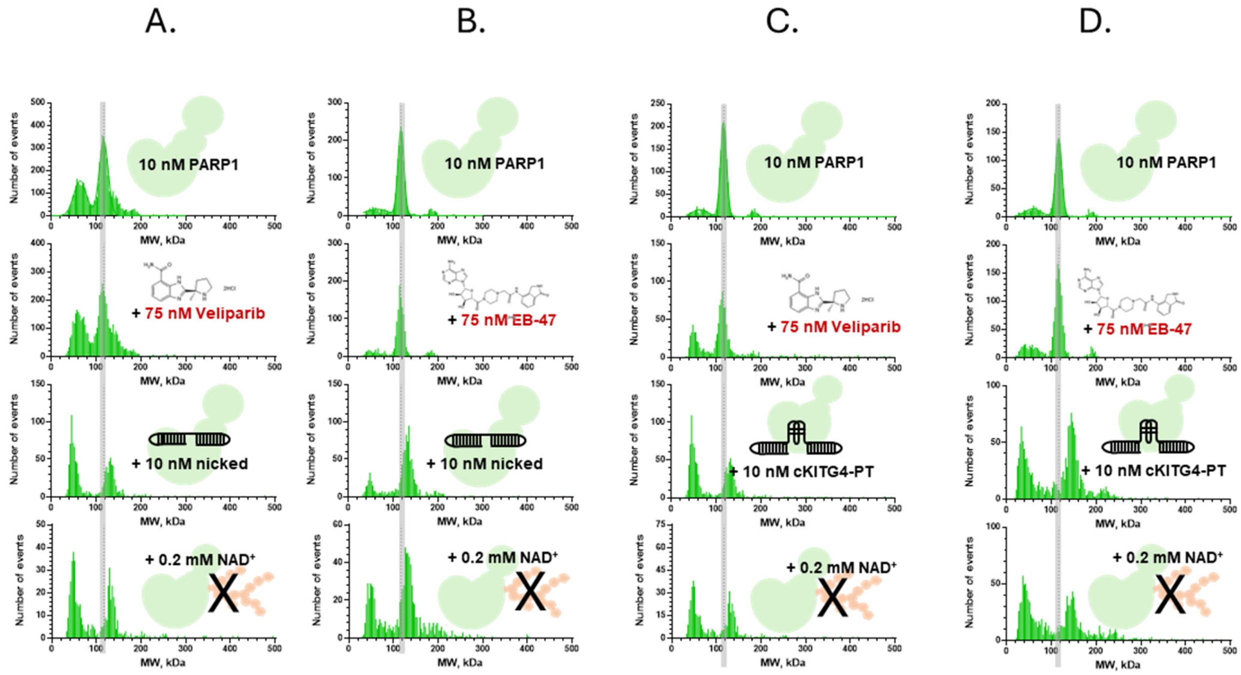


**Supplementary Figure 11.** Mass photometry analysis of DNA substrate interactions with PARP1 in presence of PARPi. Panels A-B and C-D depicts nicked and cKIT-PT DNA substrates, revealing a molecular weight shift (indicated by the transition from a gray line) upon PARP1 addition. This shift suggests the formation of PARP1-nicked and PARP1-cKITG4-PT complexes. Following the introduction of 0.2 mM $NAD^+$, there is no formation of high molecular weight PAR chains, demonstrating the PARylation process is inhibited in each case A-B (nicked) and C-D (cKITG4-PT).

`

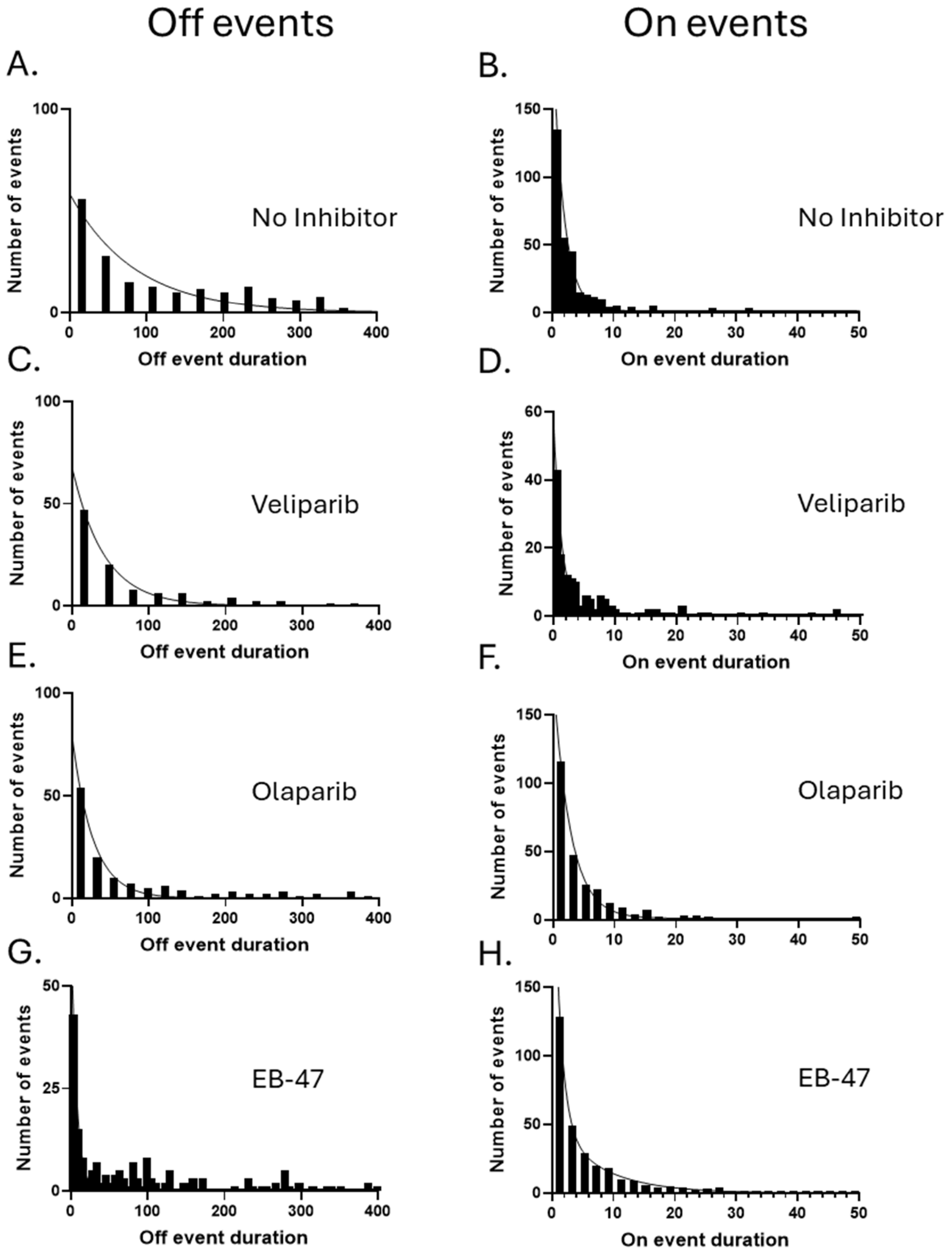


**Supplementary Figure 12.** Dwell time distributions of PARP1 on nicked DNA in the presence of PARPi. Biotinylated partial duplex DNA containing the nicked sequence was immobilized on a surface, while Cy3-labeled PARP1 (Cy3-PARP1) was infused into the reaction chamber with or without PARPi. Dwell-time distributions, constructed from all “OFF” and “ON” states, were analyzed and fitted, **(A-B)** Cy3-PARP1 in the absence of PARPi. **(C-D)** Cy3-PARP1 in the presence of Veliparib. **(E-F)** Cy3-PARP1 in the presence of Olaparib. **(G-H)** Cy3-PARP1 in the presence of EB-47.

`

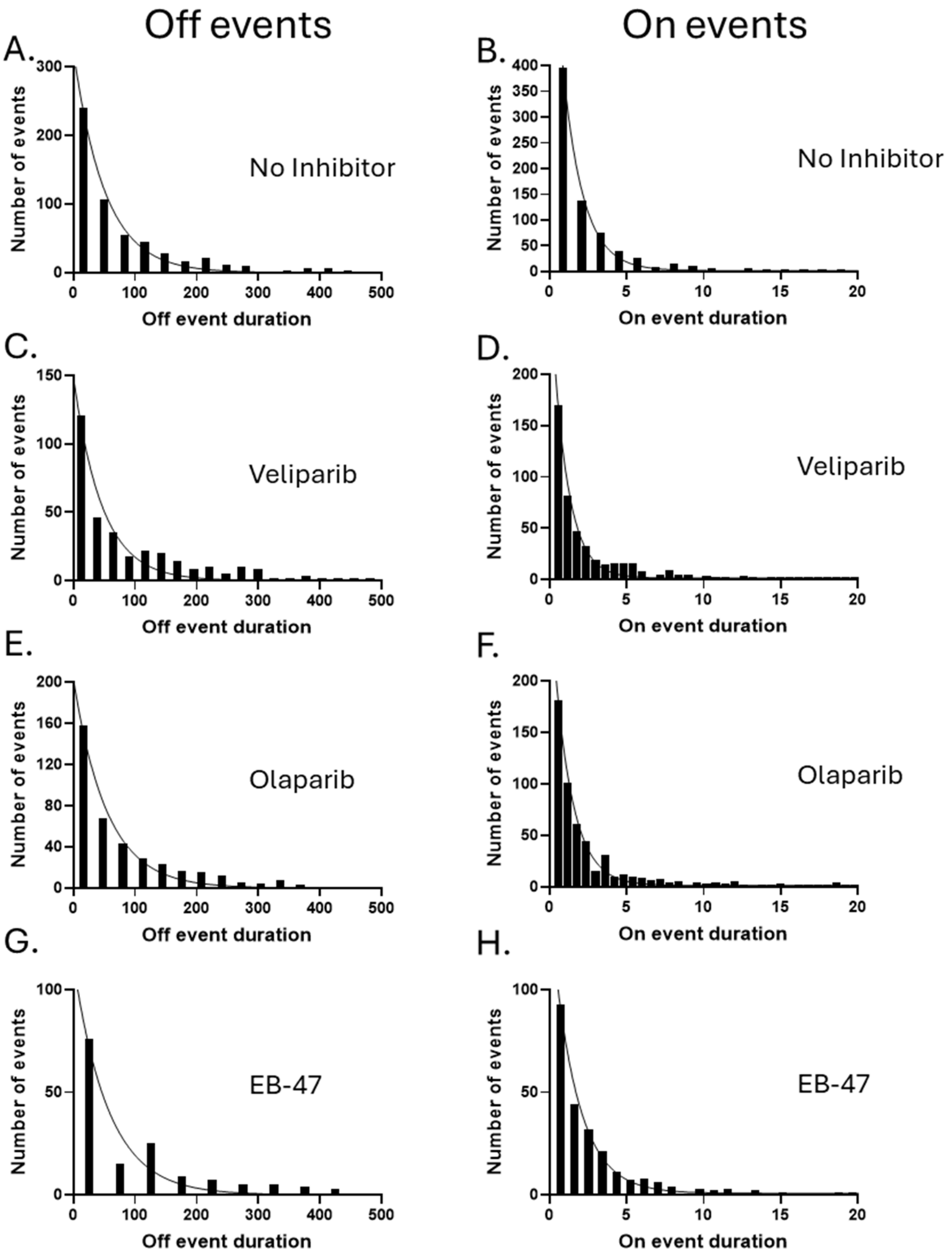


**Supplementary Figure 13.** Dwell time distributions of PARP1 on cKITG4-PT DNA in the presence of PARPi. Biotinylated partial duplex DNA containing the cKITG4-PT was immobilized on a surface, while Cy3-labeled PARP1 (Cy3-PARP1) was infused into the reaction chamber with or without PARPi. Dwell-time distributions, constructed from all “OFF” and “ON” states, were analysed and fitted. **(A-B)** Cy3-PARP1 in the absence of PARPi. **(C-D)** Cy3-PARP1 in the presence of Veliparib. **(E-F)** Cy3-PARP1 in the presence of Olaparib. **(G-H)** Cy3-PARP1 in the presence of EB-47.

`

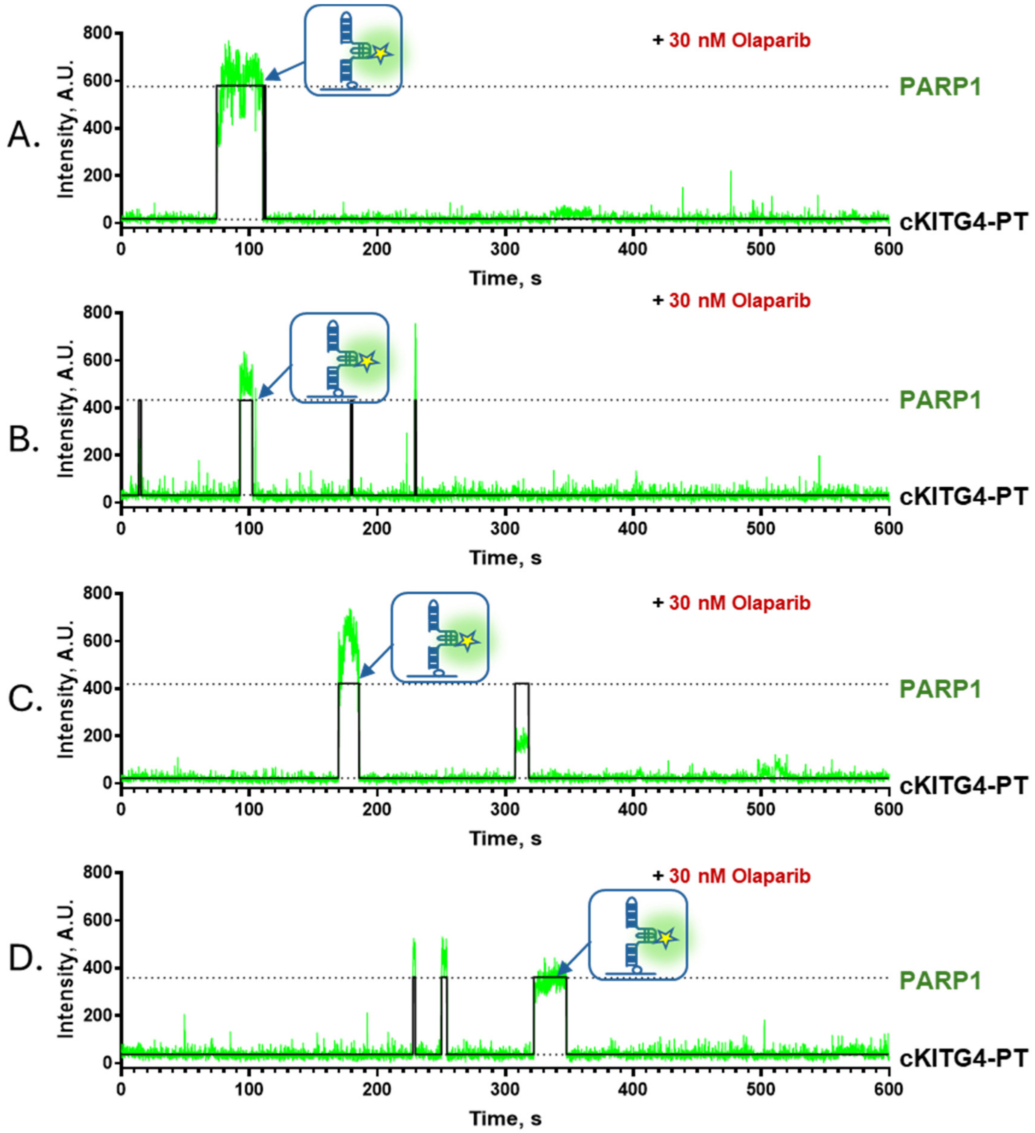


**Supplementary Figure 14.** Representative smTIRFM trajectories of Cy3-labeled PARP1 binding to surface-tethered cKITG4-PT DNA (A-C) in presence of Olaparib. Biotinylated DNA constructs were immobilized on a surface, while Cy3-PARP1 were infused into the reaction chamber. The representative trajectories show a fluorescence trajectory (green) overlaid with an idealized trajectory (black).

`

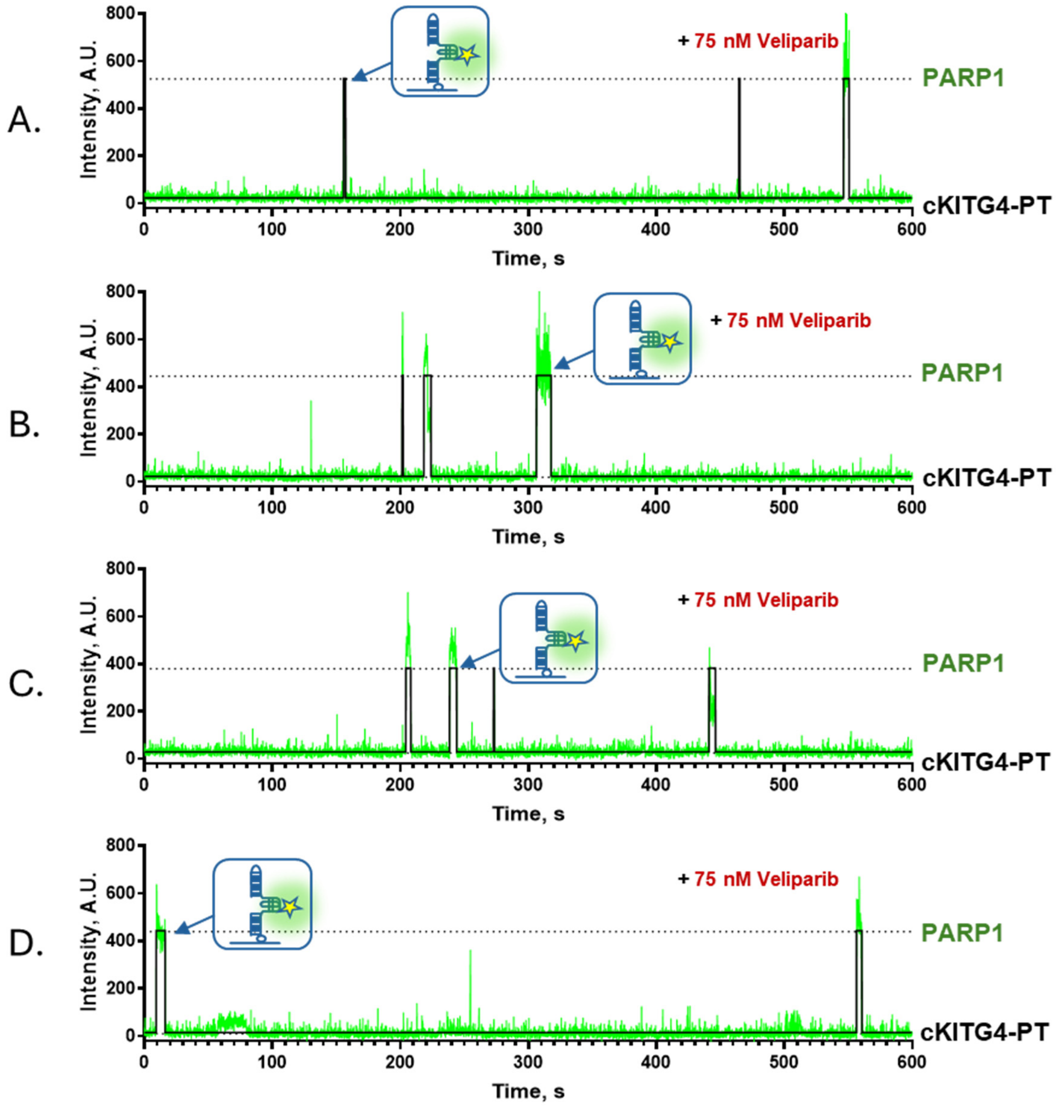


**Supplementary Figure 15.** Representative smTIRFM trajectories of Cy3-labeled PARP1 binding to surface-tethered cKITG4-PT DNA (A-D) in presence of Veliparib. Biotinylated DNA constructs were immobilized on a surface, while Cy3-PARP1 were infused into the reaction chamber. The representative trajectories show a fluorescence trajectory (green) overlaid with an idealized trajectory (black).

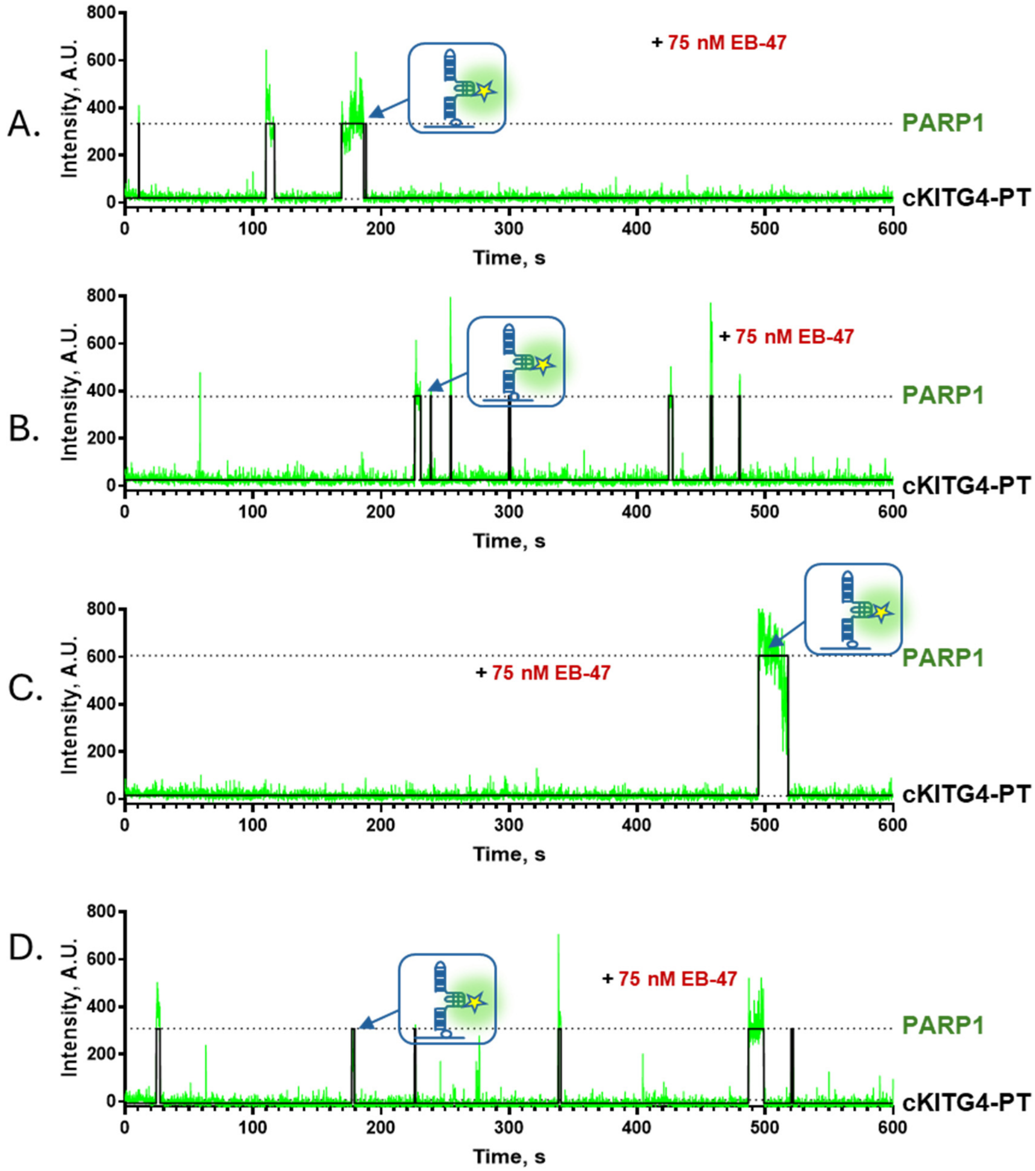


**Supplementary Figure 16.** Representative smTIRFM trajectories of Cy3-labeled PARP1 binding to surface-tethered cKITG4-PT DNA (A-E) in presence of EB-47. Biotinylated DNA constructs were immobilized on a surface, while Cy3-PARP1 were infused into the reaction chamber. The representative trajectories show a fluorescence trajectory (green) overlaid with an idealized trajectory (black).

`

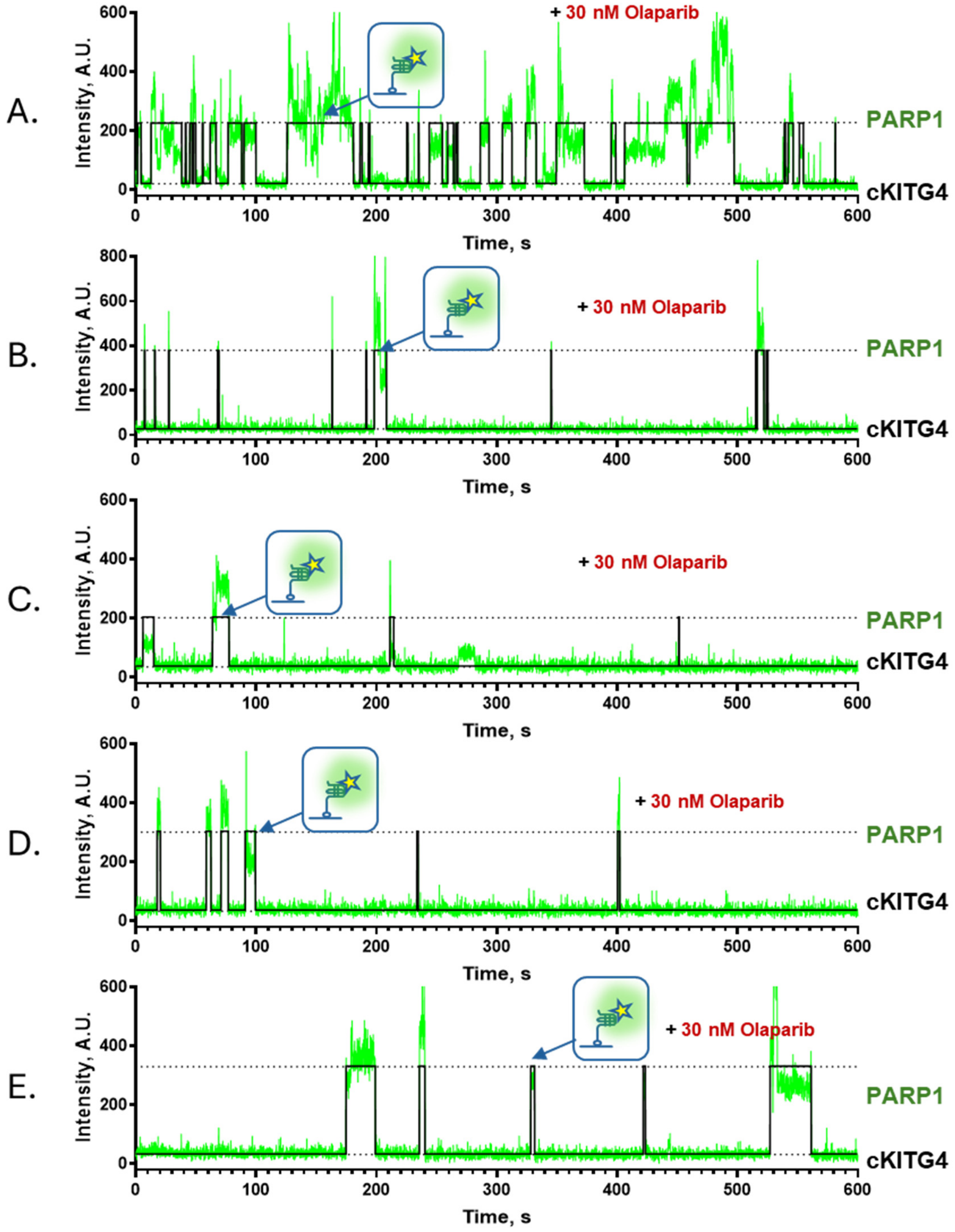


**Supplementary Figure 17.** Representative smTIRFM trajectories of Cy3-labeled PARP1 binding to surface-tethered cKITG4 DNA (A-E) in presence of Olaparib. Biotinylated DNA constructs were immobilized on a surface, while Cy3-PARP1 were infused into the reaction chamber. The representative trajectories show a fluorescence trajectory (green) overlaid with an idealized trajectory (black).

`

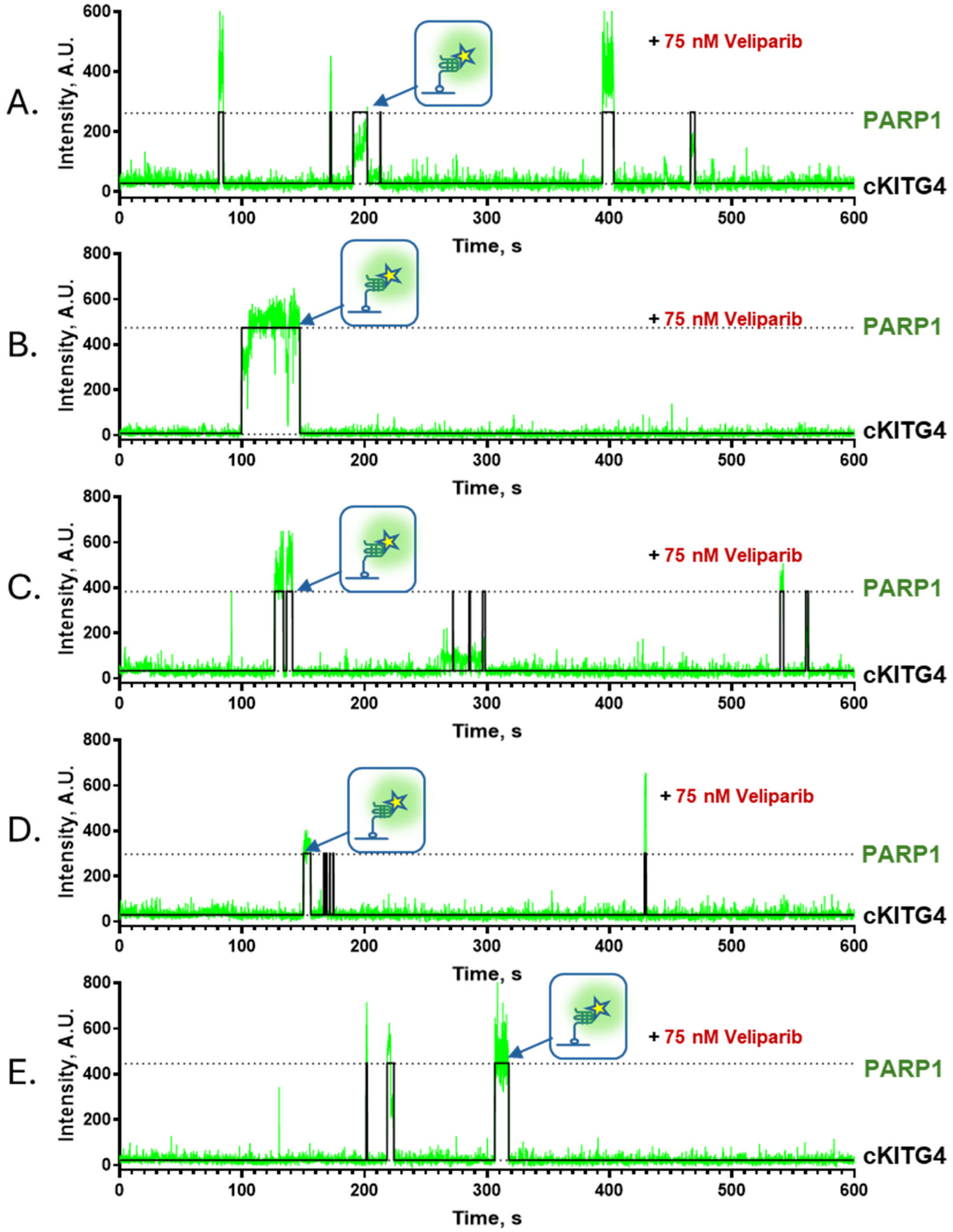


**Supplementary Figure 18.** Representative smTIRFM trajectories of Cy3-labeled PARP1 binding to surface-tethered cKITG4 DNA (A-E) in presence of Veliparib. Biotinylated DNA constructs were immobilized on a surface, while Cy3-PARP1 were infused into the reaction chamber. The representative trajectories show a fluorescence trajectory (green) overlaid with an idealized trajectory (black).

`

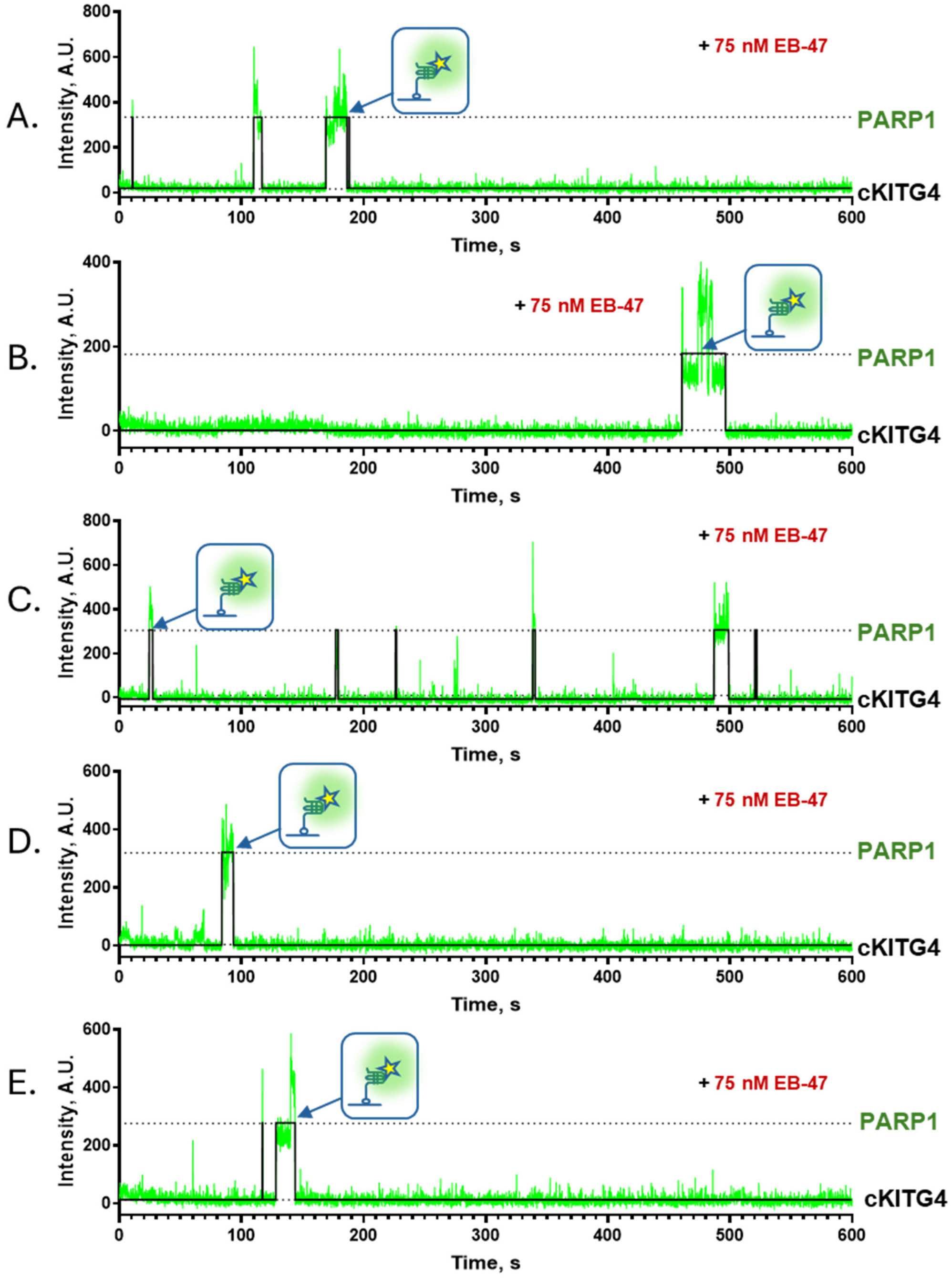


**Supplementary Figure 19.** Representative smTIRFM trajectories of Cy3-labeled PARP1 binding to surface-tethered cKITG4 DNA (A-E) in presence of EB-47. Biotinylated DNA constructs were immobilized on a surface, while Cy3-PARP1 were infused into the reaction chamber. The representative trajectories show a fluorescence trajectory (green) overlaid with an idealized trajectory (black).

`

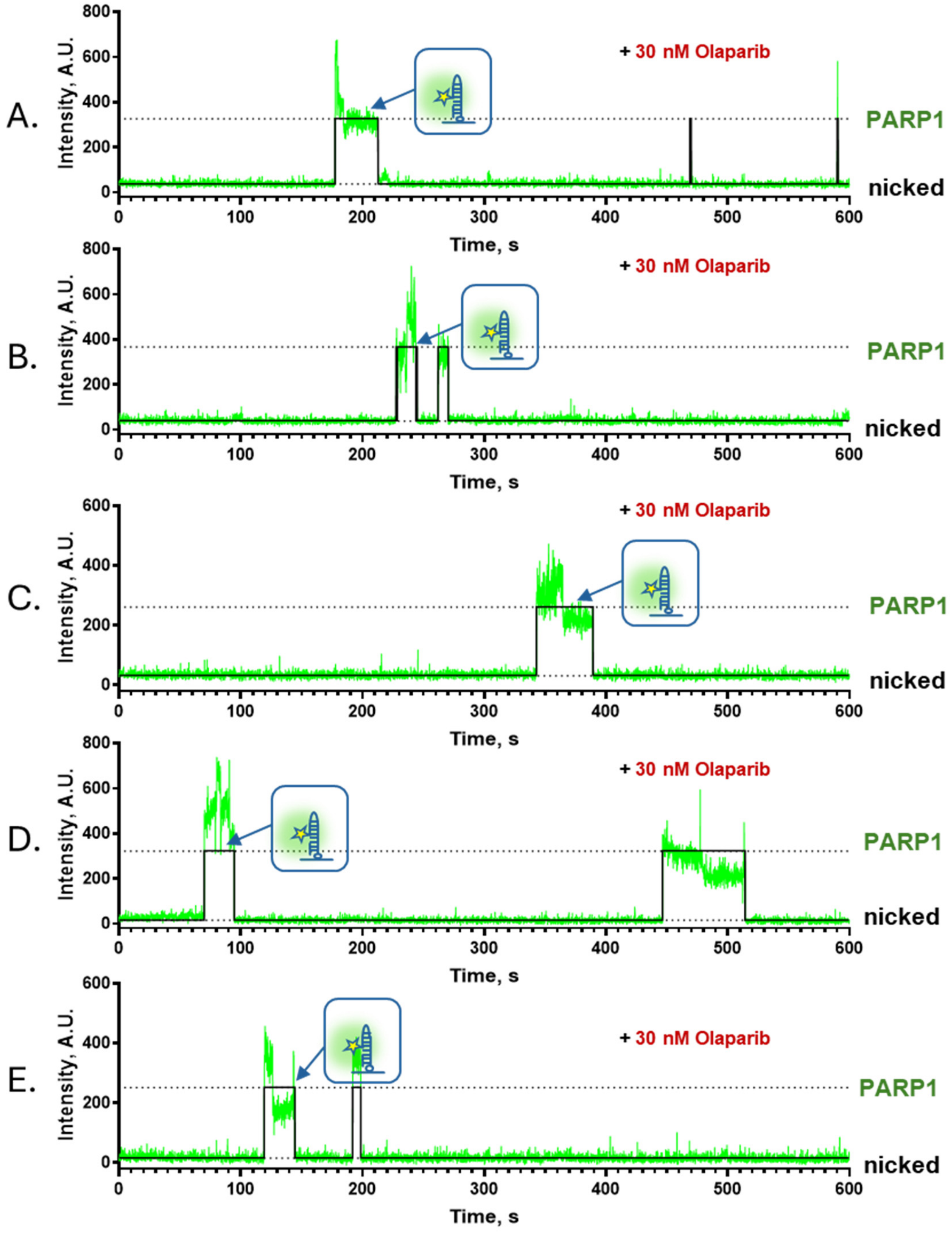


**Supplementary Figure 20.** Representative smTIRFM trajectories of Cy3-labeled PARP1 binding to surface-tethered nicked DNA (A-E) in presence of Olaparib. Biotinylated DNA constructs were immobilized on a surface, while Cy3-PARP1 were infused into the reaction chamber. The representative trajectories show a fluorescence trajectory (green) overlaid with an idealized trajectory (black).

`

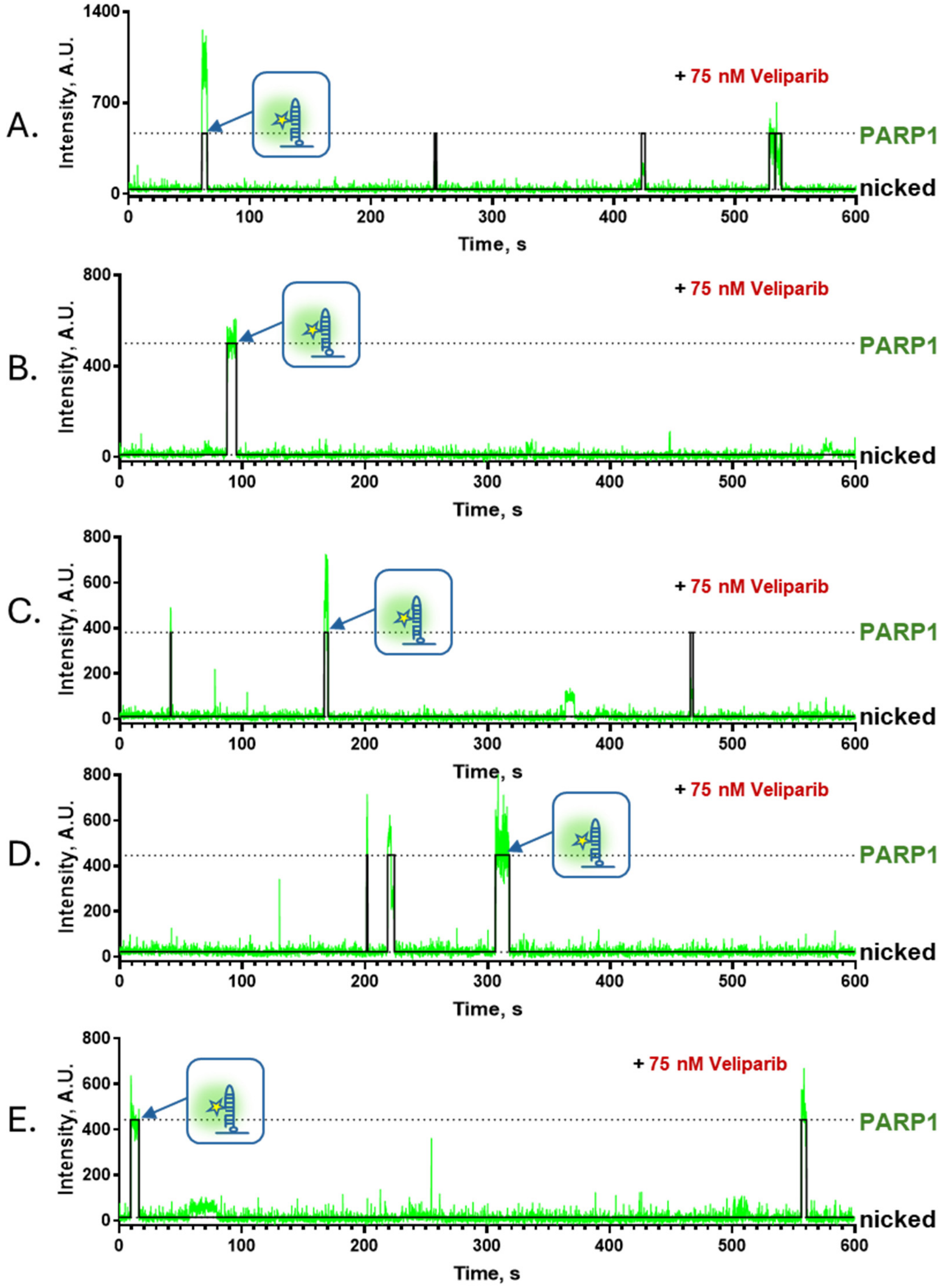


**Supplementary Figure 21.** Representative smTIRFM trajectories of Cy3-labeled PARP1 binding to surface-tethered nicked DNA (A-E) in presence of Veliparib. Biotinylated DNA constructs were immobilized on a surface, while Cy3-PARP1 were infused into the reaction chamber. The

`

representative trajectories show a fluorescence trajectory (green) overlaid with an idealized trajectory (black).

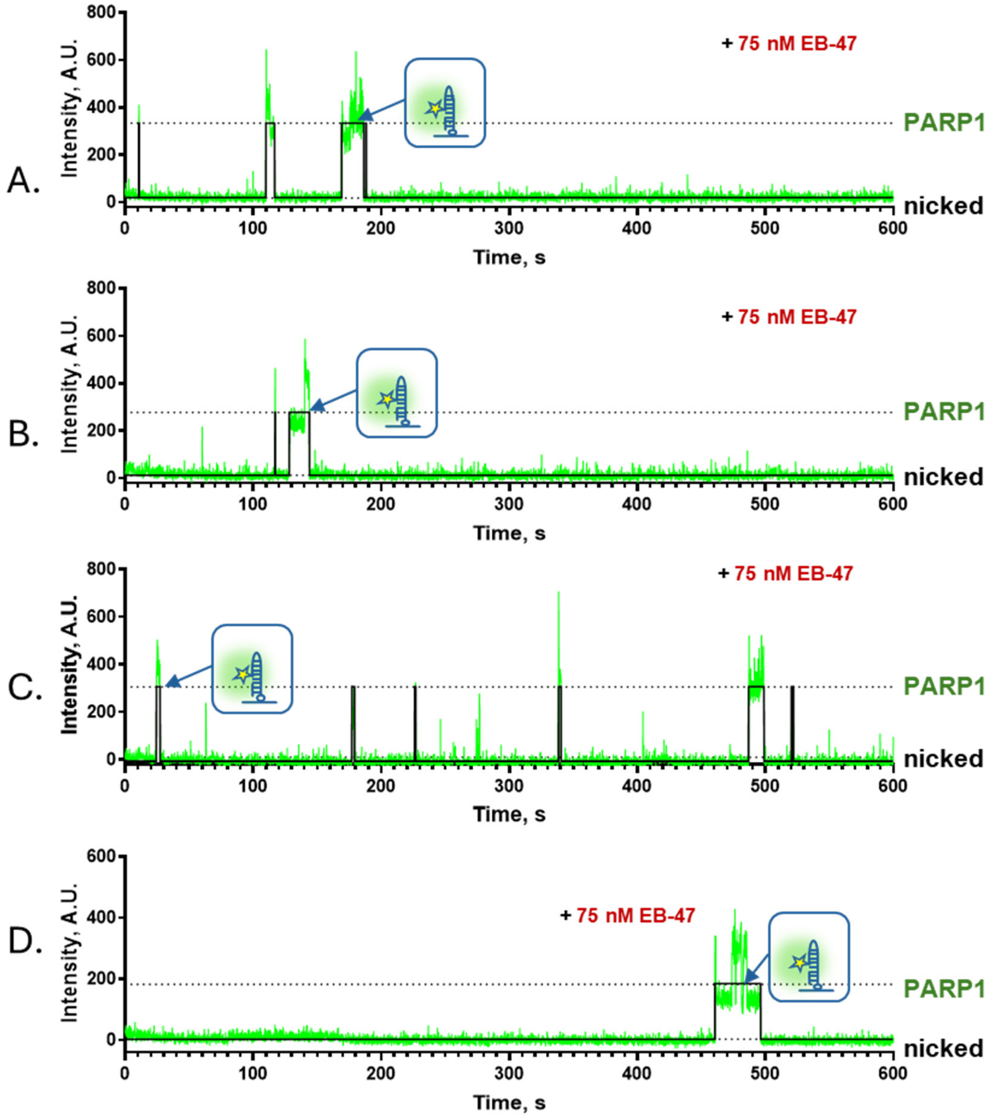


**Supplementary Figure 22.** Representative smTIRFM trajectories of Cy3-labeled PARP1 binding to surface-tethered nicked DNA (A-D) in presence of EB-47. Biotinylated DNA constructs were immobilized on a surface, while Cy3-PARP1 were infused into the reaction chamber. The representative trajectories show a fluorescence trajectory (green) overlaid with an idealized trajectory (black).

`
